# Transcriptomics and proteomics of projection neurons in a circuit linking hippocampus with dorsolateral prefrontal cortex in the human brain

**DOI:** 10.1101/2024.06.12.598714

**Authors:** Christopher Borcuk, Rahul A. Bharadwaj, Gianluca C. Kikidis, Vamshi Mallepalli, Leonardo Sportelli, Alessandro Bertolino, Robert N. Cole, Lauren R. DeVine, Brady J. Maher, Srinidhi R. Sripathy, Madhur Parihar, Joo Heon Shin, Yong Kyu Lee, Carly Montoya, Amy D. Soboslay, Thomas M. Hyde, Joel E. Kleinman, Daniel R. Weinberger, Giulio Pergola

## Abstract

Transcriptome and proteome sequencing of brain tissue homogenate has helped unravel processes underlying schizophrenia (SCZ). However, most studies have lacked granularity at the cell type level and have focused on individual brain regions, rather than examining expression dynamics across multiple regions or illness-relevant circuitries. We used laser capture microdissection to collect excitatory neuron-enriched samples from hippocampal subregions CA1 and presubiculum (SUB), and from dorsolateral prefrontal cortex (DLPFC), a circuit prominently implicated in schizophrenia. Using RNA sequencing and quantitative proteomics, we show significantly superior discrimination of brain regional identity in the transcriptomic (>90% accuracy) and proteomic data (>97% accuracy) compared with gene-level expression data (<70% in bulk). Patients with SCZ show hippocampal-specific differential protein phosphorylation. SCZ risk co-expression gene-sets that replicate across transcript and protein networks are enriched for transmembrane transporters in the DLPFC and CA1 and postsynaptic processes in the SUB. We demonstrate a strong directional connectivity effect of SCZ risk in that excitatory synaptic genes in CA1 unidirectionally predict gene expression in SUB. Finally, parallel CA1 snRNA-seq results suggest that in SCZ excitatory efferents in CA1 are affected by interactions with glia and by downregulation of inhibitory neuropeptide inputs. Our study proposes molecular mechanisms by which hippocampal communication, previously associated with SCZ at the macroscopic level, may be altered at the inter-field and interregional circuit level.

## Introduction

The latest Psychiatric Genomics Consortium (PGC) study of schizophrenia (SCZ) has identified 287 genetic loci individually conferring small effects on risk for SCZ ^1^, which were collectively proximal to genes enriched for synaptic function. An important current challenge in SCZ research is the question of risk convergence: how do hundreds of spatially distant genetic loci together confer risk for SCZ, and which biological pathways, brain regions, and circuit mechanisms do they converge upon? Studies in the postmortem human brain have associated SCZ risk with gene expression by profiling messenger RNA extracted primarily from bulk homogenate tissue. Using relatively large sample sizes, such studies have identified eQTLs and co-expression networks potentially implicated in SCZ risk ^2–10^. These insights from bulk tissue data have been further translated into potential pathophysiological mechanisms of SCZ at gene ^6,9–17^, transcript ^4,8,15^, and, most recently, protein levels ^18^.

Scientific insight is as good as the biological significance of the assays used to derive it. A major caveat of most previous studies is that information on cell specificity is mixed across cells and thus difficult to extract from tissue homogenates. Instead, single-cell and single-nuclei approaches (snRNA-seq) allow for the identification of many separate cell type clusters at once ^19–22^. However, snRNA-seq only captures nuclei rather than whole cell bodies. Therefore, the synthesized mRNA transported to peripheral cell components, such as the synapse, cannot be reliably quantified in the nucleus, with decreased RNA yield compared to whole cells. Further, cell types are inferred only by their genetic profile, while morphological or other anatomical features are not considered; finally, 3’ amplification bias, a limitation of the most widely used snRNA-seq approach, limits the potential for isoform resolution ^23^. These caveats can be addressed with the use of laser capture microdissection (LCM), a significantly more cost-effective approach that enables the selection and isolation of whole cells prior to RNA-seq or proteome analysis ^24^. LCM also complements snRNA-seq, given its morphology-dependent cell population selectivity.

We have previously used LCM to collect cell-type-enriched data from the dentate gyrus of 263 subjects ^3,6^. Unsurprisingly, DG-LCM data showed a greater proportion of excitatory neurons and greater expression of neuronal genes as compared to bulk-derived data from the hippocampus (HP) of the same individuals. Fifteen DG-specific SCZ risk eQTLs undetected in bulk hippocampal tissue were identified in the LCM-enriched cell samples. Co-expression networks derived from DG-LCM were more faithful to neuronal gene ontologies and presented a greater SCZ risk gene enrichment than bulk tissue ^6^. The granule cell layer of the DG is nonetheless quite compact and easy to capture morphologically, and an LCM analysis targeting interregionally distributed brain cell populations classically linked to SCZ has yet to be tested.

Here, we have utilized LCM to investigate the correspondence of gene expression based on RNA-seq and individual proteins measured by quantitative proteomics using Tandem Mass Tags (TMT proteomics) in a classical circuit implicated in hippocampal-prefrontal connectivity and comprised of hippocampal-prefrontal pyramidal neurons. We hypothesized that differences between cases and controls would be apparent at the circuit-level, even if not detectable when considering one region at a time. Specifically, we expected to find evidence of abnormal HP-DLPFC transcriptomic coherence, mirroring prior magnetic resonance imaging evidence of abnormal connectivity between these brain regions ^25^.

We used LCM to collect cell-specific RNA-seq and TMT proteomics data from 10 patients with SCZ and 10 demographically matched healthy controls (CTR) in a circuit connecting the hippocampal formation subregions CA1 and presubiculum (SUB) with DLPFC, long implicated in SCZ pathogenesis ^26^ (Figure 1A). We isolated projection neurons in mono-synaptic and reciprocally connected regions ^27,28^, i.e., from glutamatergic neurons in layer III of DLPFC ^7,12,29,30^, in SUB, and CA1 of the HP ^25,31,32^ (Figure 1B). In parallel, we performed snRNA-seq in the CA1 to assess how cell type interactions in one brain region may influence circuit dynamics.

**Figure 1.**
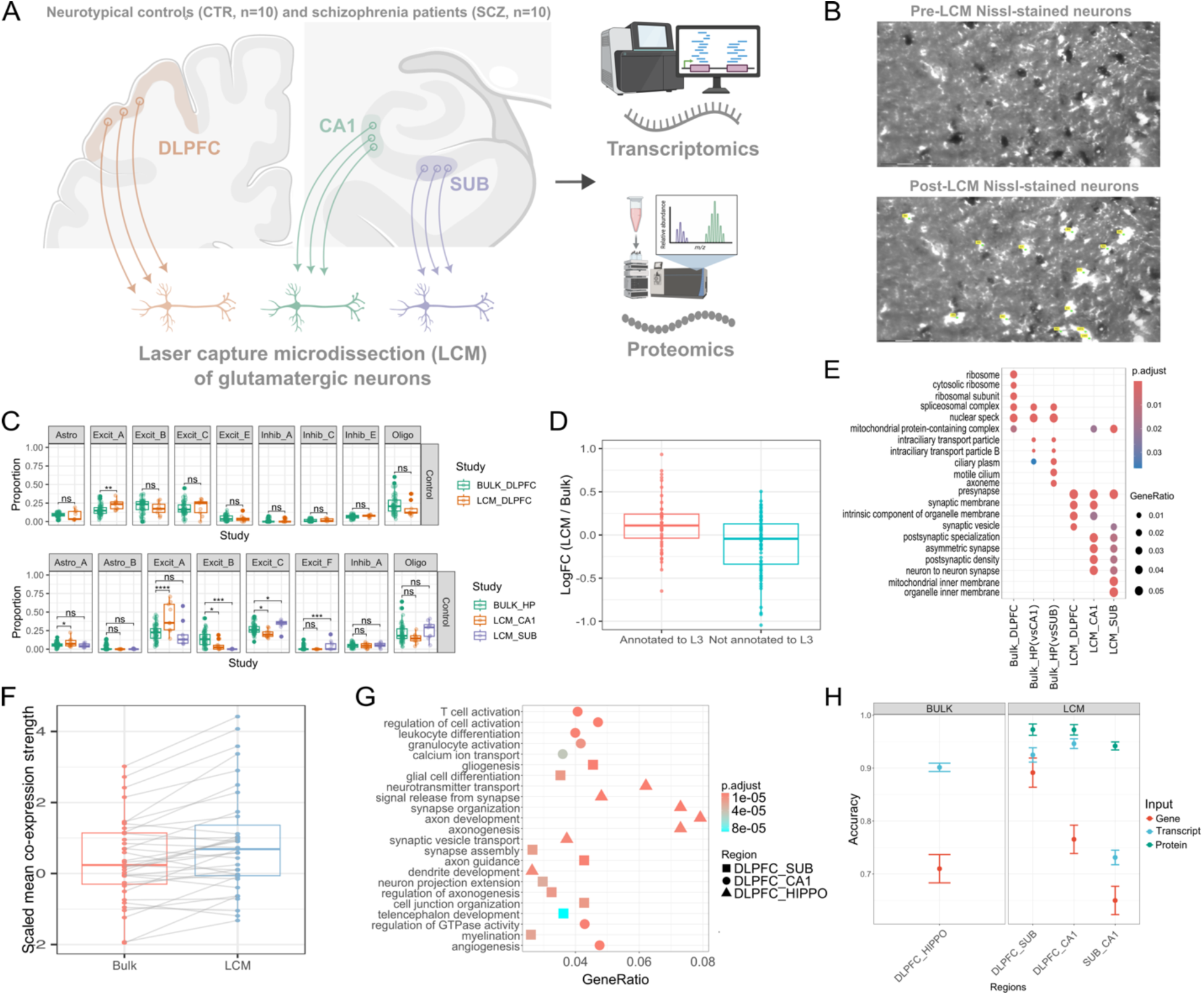
Comparison of gene expression in bulk vs LCM tissue. **A)** Major projection neurons between the hippocampal output centers CA1 and SUB (bottom left), and DLPFC (top left) were extracted using Laser Capture Microdissection (LCM), then sequenced at the transcriptome and proteome levels. (CA1 - cornu ammonis stratum pyramidale; SUB- subiculum large pyramidal layer, DLPFC - dorsolateral prefrontal cortex). **B)** Sequential isolation (selection, cutting, and visualization) of Nissl-stained projection neurons (representative example is from CA1 stratum pyramidal neurons) using Zeiss Laser Capture Microdissection (LCM). **C)** Cell type proportions were estimated using CIBERSORTx in hippocampal tissues **(bottom panels)** and DLPFC **(top panels).** We used DLPFC and HP snRNA-seq data from Tran et al., 2021 as a reference ^20^. LCM tissues see the specific increase in proportion of certain excitatory neuronal subtypes (from HP snRNA signature matrix CA1 = Excit_A, SUB = Excit_F, and Excit C; from DLPFC snRNA signature matrix DLPFC = Excit_A) as compared to bulk tissue. **D)** We evaluated if marker genes for L3 were more differentially expressed in LCM versus Bulk as compared to marker genes for other layers. We used markers published from a spatial transcriptomics analysis of the DLPFC ^35^. **E)** We checked Gene Ontology (cellular compartment) enrichments for significantly differentially expressed genes (DEGs) between Bulk and LCM samples. The y-axis shows enriched terms for up-regulated DEGs in each brain region (x-axis). **F)** Comparison of within-set scaled mean co-expression strength values of neuronal CLIC gene sets between LCM and bulk tissue. **G)** Machine learning prediction of regional identity. We established differentially expressed genes between each brain region to assess regional identity through a random forest leave-one-out cross-validation analysis. The gene ontology enrichment for the most predictive genes. **H)** shows average prediction accuracy and error bars between DLPFC/HIPPO (Bulk tissue) and DLPFC/SUB, DLPFC/CA1, and SUB/CA1 (LCM data). Red dots represent gene-level data, blue dots represent transcript data, and green dots represent protein data. Abbreviations: LCM, laser capture microdissection; SCZ, schizophrenia; CTR, control.

Our TMT proteomics data identifies differential peptide phosphorylation in SCZ, especially in the HP, implicating cytoskeletal processes. By focusing on transcriptomic findings that are mirrored in proteomic expression, we highlight genes that show high translatability; these are shown to be strongly neuronal and with relative transcriptional stability over time, as assessed via longitudinal sampling in human iPSC-derived neurons. We compute gene co-expression networks in transcriptomic and proteomic states and identify replicable SCZ co-expression partners across both single-cell methods. Using novel interregional co-expression techniques, we show that CA1 genes related to SCZ risk are associated with excitatory synaptic function downstream in the SUB rather than the reverse, implicating a directional mechanism of risk-associated cell-cell interaction. Finally, snRNA-seq shows CA1 excitatory clusters to have reduced inputs from inhibitory cells and increased communication with glial cells, showcasing potential modulatory elements influencing CA1 excitatory efferents.

## Results

### LCM-derived data compared with bulk tissue

To test the biological granularity of our LCM data, we first compared the quality of LCM pyramidal neuron enrichment data to bulk tissue RNA-seq from the Lieber Institute for Brain Development (LIBD) brain repository ^5^. We verified that gene expression differed in cell proportions obtained via deconvolution approaches compared with bulk tissue (CIBERSORTx with published 10x DLPFC and HP data by Tran et al. ^20^). LCM samples showed a relatively increased proportion of specific excitatory neuronal subtype clusters (Figure 1C, subtype comparisons; Tukey p <.001). The LCM CA1 was enriched for the HP Excit A snRNA cell type cluster, which prior reports consistently ascribed to CA1 ^20,33^. The LCM SUB was enriched for Excit D and Excit F, indicating, as expected, an enrichment of pyramidal neurons over granule cells. The LCM DLPFC was enriched for the DLPFC Excit A snRNA cell type cluster, again consistent with prior reports on middle (L3-L4) DLPFC layers^20,34^. We evaluated how genes were differentially expressed between LCM and Bulk tissues and found that marker genes for layer III of DLPFC, according to a spatial transcriptomics study of this region ^20^, were more strongly expressed in LCM DLPFC compared with bulk (Figure 1D). Genes with significantly increased differential expression in LCM compared with bulk showed consistent enrichment for synaptic ontologies (Figure 1E). Together, these results showcase the increased neuronal expression signal obtained in LCM enriched samples as compared to bulk tissue.

### LCM co-expression profiles show increased co-expression strength in neuronal gene sets

Gene co-expression techniques are often used for parsing out molecular pathways present in expression data. We hypothesized that LCM data would have superior resolution to define the co-expression of synaptic and neuronal gene sets compared with bulk tissue data. We used the CLustering by Inferred Co-expression (CLIC) database to extract gene sets consistently co-expressed across published RNA expression datasets. CLIC sets were clustered into communities based on gene overlap (Figure S3A), and a neuronal community was identified based on overrepresentation of neuronal GO terms in overlapping genes (Figure S3B) and overrepresentation of neuronally relevant words in CLIC set names (Figure S3C). We used a permutation approach to evaluate if a gene set is more co-expressed with itself than the background and found that the relative co-expression strength of neuronal gene sets was significantly increased in LCM as compared with bulk (Figure 1F, t(18) = 4.4, p <.001). This evidence indicates that LCM neuron-enriched samples have a greater potential to precisely parse neuronally relevant pathways related to neuropsychiatric risk.

### Machine learning prediction of regional identity

To assess the biological information content of the different omics assays we collected, we predicted brain region (hippocampus or DLPFC) based on gene expression at the gene, transcript, and proteomic levels. We determined differentially expressed genes between regions in a leave-one-out cross-validation framework and used a Random Forest classifier to predict the region of the left-out sample. We computed 100 iterations to generate reliable distributions of expression profiles between brain regions. We then compared prediction accuracies within the same samples between the different omics data.

Figure 1G shows the gene ontology enrichment for the most predictive genes for each region pair comparison. Figure 1H shows the prediction results. At the total gene level, bulk gene-level reads distinguish DLPFC/HP at 69% accuracy, while LCM mRNA improves DLPFC/SUB and DLPFC/CA1 predictions to 88% and 72.5%, respectively. At the transcript level, LCM yields 92.5% (DLPFC/SUB) and 94.6% (DLPFC/CA1) accuracy, outperforming bulk (DLPFC/HP: 84.2%). These LCM RNA-seq improvements over bulk were significant (gene: DLPFC/HP vs DLPFC/SUB, *t*(79)=5.53, *p*<.001; vs DLPFC/CA1, *t*(79)=3.83, *p*<.01; transcript: DLPFC/HP vs DLPFC/SUB, *t*(79)=5.50, *p*<.01; vs DLPFC/CA1, *t*(79)=6.8, *p*<.001). LCM TMT proteomics, however, achieved the highest accuracy at 97.5% (DLPFC/SUB) and 97.4% (DLPFC/CA1), significantly outperforming LCM transcript predictions (DLPFC/SUB: *t*(39)=44.6, *p*<.001; DLPFC/CA1: *t*(38)=11.5, *p*<.001). These results confirm the expectation that brain region identity is better represented in our LCM samples as compared to bulk, and additionally reveal that the transcript and proteomic level in LCM outperform gene-level data.

### Transcriptome to proteome translatability and its relation to mRNA stability in iPSC-derived neurons

As transcriptomic and proteomic evaluations of expression have been found divergent in past research^36,37^, we aimed to identify genes consistently expressed between transcriptomic and proteomic states and characterize them through gene ontology enrichment analysis. We computed a “translatability score” per gene as the across-subject association between RNA-seq and TMT proteomics data. We identified separate sets of genes with nominal significance for the linear RNA-protein association for each brain region and assessed gene ontology enrichments. Genes with significant RNA-protein translatability, accounting for 13% of gene/protein pairs in the CA1 and DLPFC and 23% of pairs in the SUB, were most strongly enriched for synaptic terms across both hippocampal tissues and for cell surface in the DLPFC (Figure 2A).

**Figure 2.**
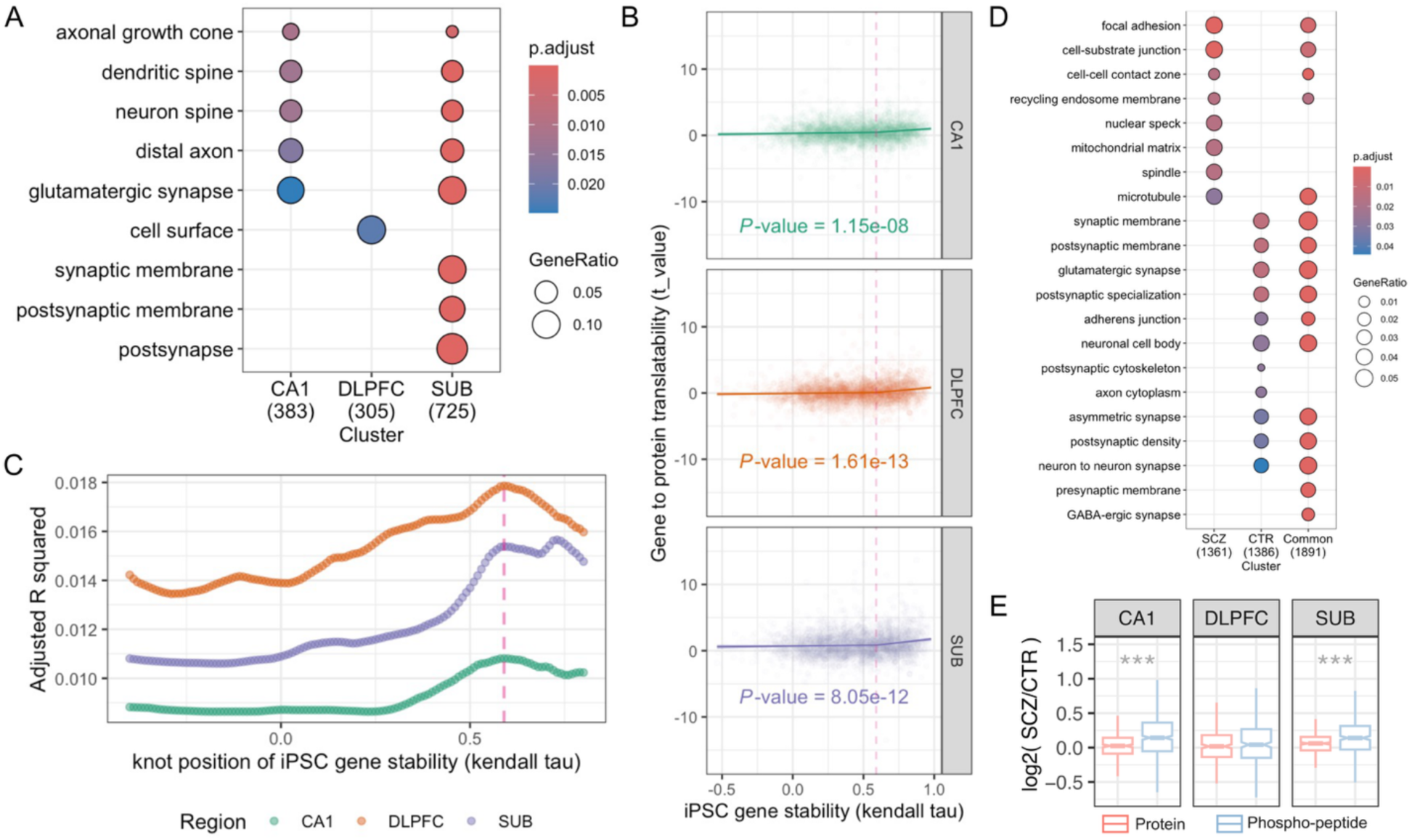
Transcriptome to proteome translatability and its relation to iPSC mRNA expression stability. **A)** The strongest gene ontology terms (biological process) enriched in sets of genes whose expression profiles did not differ between transcriptomic and proteomic states. The color scale depicts the adjusted p-value of the ontology enrichment, and the size of the dot depicts the ratio of gene ontology genes in the set. **B)** The relationship between the translatability of a gene between transcriptomic and proteomic states and the stability of a gene in iPSCs. On the y-axis, “Gene to protein translatability” is computed as the t-value arising from a linear model between gene expression and protein expression across subjects, including relevant confounders. On the x-axis, iPSC gene stability is computed as the Kendall tau correlation coefficient between DIV56 expression and DIV70 expression across iPSC samples. Points represent genes. The line represents the best-fit line based on a spline function with a knot at iPSC gene stability = 0.59. **C)** This knot value was shown to result in the highest adjusted R2 for the CA1 and DLPFC and was the second highest local maxima in SUB after knot = 0.72. The y-axis shows the adjusted R2 value when using a specific knot value for iPSC gene stability (found on the x-axis) for the spline regression of “Gene to protein translatability” vs. “iPSC gene stability.” **D)** Enrichment for genes found significantly stable from DIV52 to DIV70 (Kendall fdr adjusted p < 0.05) using only CTR subjects or only SCZ subjects. Genes found uniquely significant using only CTR subjects-derived stability are labeled “CTR,” genes found uniquely significant using only SCZ subjects-derived stability are labeled “SCZ,” and genes with significant stability in both are labeled “Common.” **E)** SCZ-dependent log fold change values (log2(SCZ/CTR)) were computed for each protein and their corresponding phospho-peptides. Only proteins with corresponding phospho-peptides were included. Values of log2(SCZ/CTR) were compared between proteins and phospho-peptides using a Wilcoxon rank sum test within each region. ***: p <.001. Abbreviations: DIV. days in vitro; iPSC, induced pluripotent stem cells; SCZ, schizophrenia; CTR, control.

Given that transcription and translation vary over time depending on cellular states and activity, we hypothesized that stronger translatability would be related to increased gene expression stability over time. To assign a stability measure per gene, we used published iPSC-derived neuron RNA-seq expression data ^38^. We considered a gene to have stronger stability if it had similar expression profiles between two mature neuronal states, occurring at days in vitro (DIV) 56 and 70 in this case. The iPSC gene stability measure was thus computed as the across-subjects Kendall’s tau correlation coefficient between gene expression at DIV56 vs DIV70. Across all tissues, the relationship between gene translatability and stability (Figure 2B) was accurately described using a spline regression with a knot at iPSC gene stability = 0.59 (CA1 F(2,32) = 18.3, DLPFC F(2,32) = 29.7, SUB F(2,32) = 25.8, all adjusted R^2^ > 0.01,_all p <.001; Figure 2C).

Figure 2B shows no significant relationship between transcriptomics and proteomics quantifications with low stability in iPSCs (<.59 iPSC stability; CA1 F(1,21) = 2.41, DLPFC F(1,21) = 1.95, SUB F(1,21) = 1.20, all p >.1), and a strong linear relationship for high-stability genes (>.59 iPSC stability; CA1 F(1,11) = 8.87, p =.002, DLPFC F(1,11) = 15.1, p <.001, SUB F(1,11) = 14.1, p <.001).

To evaluate diagnosis-dependent effects, iPSC gene stability was also computed using iPSC-derived neuronal samples only from SCZ or only from CTR subjects, and gene ontology cellular compartment enrichment was evaluated for significantly stable genes (Figure 2D, genes with iPSC stability >0.59 FDR-adjusted p <.05). Genes found significant using both SCZ- and CTR-derived stability measures were enriched for synaptic and cell adhesion terms. Genes uniquely significant to CTR-derived stability were again enriched for synaptic terms. In contrast, those uniquely significant to SCZ-derived stability were not enriched for synaptic terms but rather for cell adhesion terms, with additional unique enrichments in nuclear processes (spindle, nuclear speck) and mitochondrial matrix.

In summary, we found a positive relationship between gene stability in iPSC-derived neurons and gene-protein translatability, specifically across highly stable genes. These results suggest that previously reported low replication between transcriptomics and proteomics partly depends on the stability of assessments over time ^36,37^. By prioritizing findings that are consistent between transcriptomic and proteomic states, we may obtain more precise identification of synaptic mechanisms reliably associated with SCZ.

### Protein phosphorylation in SCZ

Protein phosphorylation is a critical posttranslational modification impacting the trafficking and activity of proteins. Altered protein phosphorylation has been observed in patients with SCZ and may contribute to the disruption of molecular pathways related to its pathophysiology ^39,40^. We thus evaluated whether SCZ-dependent changes in phosphorylation could also be observed in our TMT proteomics data. A log2(SCZ/CTR) SCZ-dependent abundance ratio value was computed for each protein and its corresponding phospho-peptides, detailing how much that protein or phospho-peptide is differentially expressed in SCZ patients compared with CTR subjects. The log(SCZ/CTR) values were compared between proteins and their phospho-peptides using a Wilcoxon rank sum test within each region. Phosphorylated peptides were shown to have higher log2(SCZ/CTR) values as compared to proteins in the CA1 and SUB (p <.001) but not in DLPFC, preliminarily suggesting hippocampal specificity to increased protein phosphorylation in SCZ (Figure 2E). Proteins and phospho-peptides with nominally significant differential expression are depicted in volcano plots in Figure S1. Gene set enrichment analysis revealed cell compartment ontologies with significantly increased or decreased SCZ differential expression, where the CA1 showed significantly decreased log2(SCZ/CTR) of proteins found in the synapse, and instead an increased log2(SCZ/CTR) of phospho-peptides found in the nucleoplasm and nuclear lumen (Figure S2).

### Co-expression networks identify SCZ risk pathways consistent across transcriptome and proteome

Using weighted gene correlation network analysis (WGCNA), we sought to identify SCZ risk enriched gene-sets that are replicable across both transcriptomic and proteomic domains (Figure 3A). Networks were computed in each region using gene, transcript, and protein data separately. For each network, we prioritized modules that were enriched for SCZ risk, using a method similar to that implemented by Pergola et al., 2023 ^6^, shown to robustly re-identify previously prioritized SCZ modules from published data. We computed overrepresentation statistics per module for six separate lists of SCZ risk genes, as well as enrichment for SCZ risk using the Multi-marker Analysis of GenoMic Annotation (MAGMA) tool ^41^, leaving seven separate SCZ risk enrichment measures per module. If a module was significantly enriched in at least three of these measures (Figure 3B, green tiles), we labeled it a SCZ risk module.

**Figure 3.**
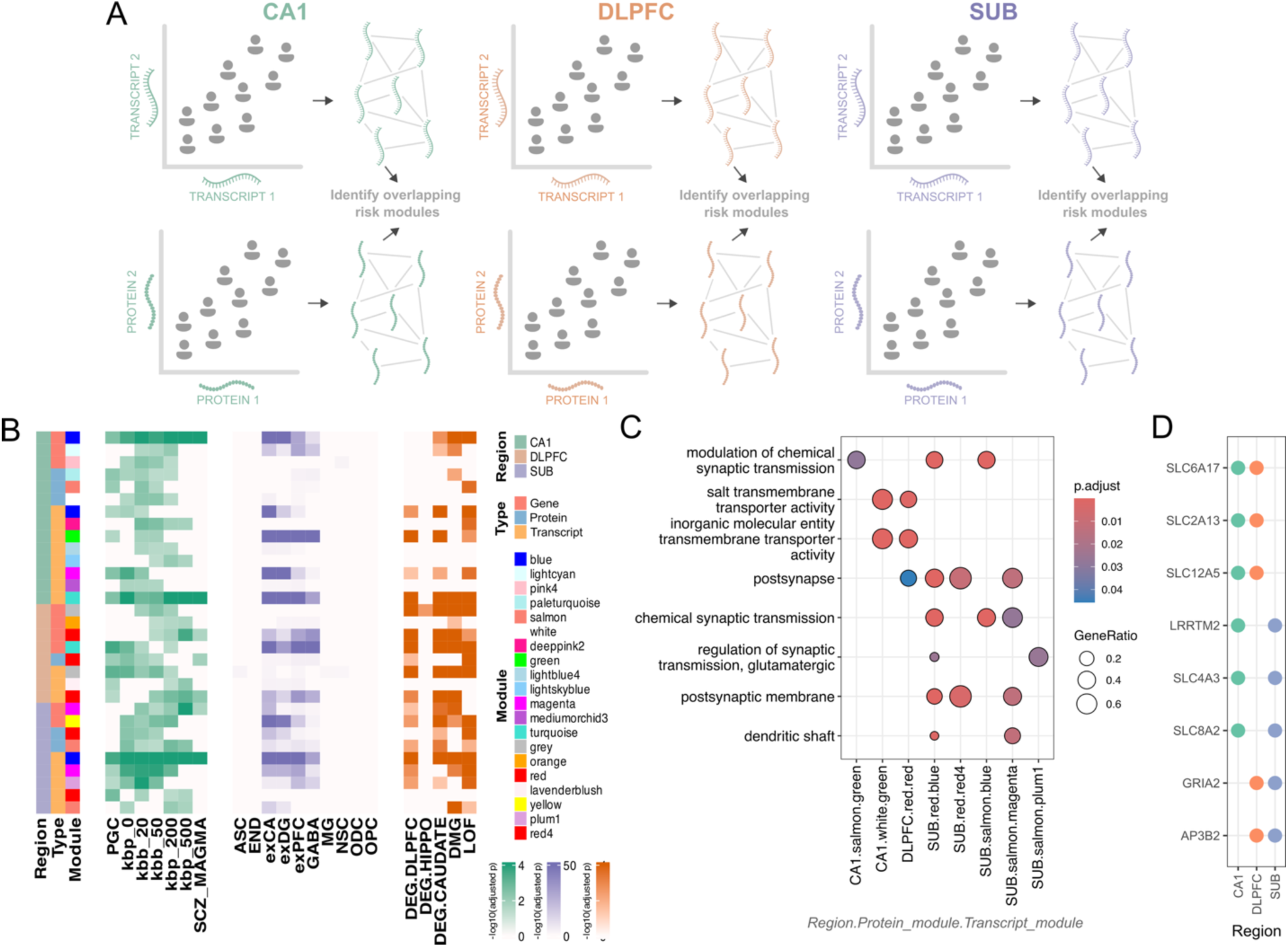
Gene co-expression and identification of replicable SCZ risk co-expression partners. **A)** WGCNA co-expression networks were constructed using gene, transcript, and protein level data. SCZ risk modules that are replicated across transcript and protein networks were prioritized. **B)** Gene enrichment analysis results for modules prioritized by SCZ risk and cross-expression type replicability. Green grids show enrichment results for SCZ risk genes and SCZ MAGMA. Purple grids show enrichment results for cell type specificity using a human single-cell atlas. Orange grids show enrichment testing results for differentially expressed genes in SCZ-CTR contrasts, differentially methylated genes, and loss-of-function variant-intolerant genes in the green grid. **C)** Gene ontologies of each significant intersection between transcript and protein prioritized modules. In parentheses are the number of genes present in the intersection. The FDR-adjusted p value is depicted with a color scale, and the ratio of present ontology genes is depicted by the size of the dot. **D)** Genes that were found in transcript and protein prioritized modules in at least 2 different regions. Abbreviations: LCM, laser capture microdissection.

We first looked at modules that had significant overlap across at least two expression measures (among gene, transcript, or protein) within the same region, using the Jaccard index (JI). For example, a SCZ risk gene module in DLPFC was only considered further if it had a significant JI with another SCZ risk module in the transcript network or the protein network of DLPFC. These modules were generally enriched for excitatory cell types matching the region of origin (Figure 3B, purple tiles), and for published differentially expressed genes between SCZ patients and CTRs (DEGs) found in the DLPFC and Caudate ^4,5^, for differentially methylated genes, and for loss-of-function genes (Figure 3B, orange tiles).

We chose to further prioritize genes found in modules that are replicated specifically across transcript and protein networks (significant JI). This procedure served to profile only the most robustly co-expressed proteins, as well as to parse the protein modules further into more specific isoform resolutions that are better resolved using our transcript isoform modules. We identified consistent synaptic ontologies related across all regions (Figure 3C). Specific to the SUB were post-synaptic glutamatergic synapse ontologies. DLPFC and CA1 showed related ontologies concerning transmembrane transporter activity. Eight genes were replicated across transcript and protein networks in at least 2 regions (Figure 3D): *SLC12A5*, *SLC2A13*, *and SLC6A17* in CA1 and DLPFC; *LRRTM2*, *SLC4A3*, and *SLC8A2* between CA1 and SUB; and *AP3B2* and *GRIA2* between DLPFC and SUB. Five genes, all found in the CA1 overlapping with either DLPFC or SUB, are of the solute carrier class (SLC), responsible for the transmembrane movement of nutrients and ions ^42^. Two genes, *GRIA2* and *LRRTM2*, both found in SUB, are related to AMPA receptor function ^43–46^. In summary, our comprehensive co-expression network analysis identifies transcript-to-protein translatable SCZ-risk gene partners that imply region-specific SCZ-dependent effects on transmembrane transport and AMPA receptor-related processes.

### Interregional circuit-level effects of SCZ diagnosis

Interregional functional networks are often shown to be perturbed in SCZ, in the context of neuroimaging studies^25,31,47^, and functional connectivity has been shown to associate with interregional gene correlation strength^48^. We, therefore, hypothesized that interregional co-ordination of specific gene sets would be affected in patients with SCZ. We measured the across-gene correlation strength between pairs of regions for each CLIC gene set and for each individual and assessed associations with diagnosis status (Figure 4A). For a specific gene set in a specific subject for a specific region pair, we computed an interregional *transcriptomic coupling* index as the across-gene Pearson R coefficient. We first evaluated transcriptomic coupling using the union of all CLIC genes per subject. This global measure was highly significant for all region pairs in all subjects (all Pearson R >.78, all p <.001) (Figure S4A), suggesting consistent ranking of gene expression between regions across all subjects. Two-way ANOVA aimed to interrogate the combined effects of region-pair and diagnosis on transcriptomic coupling, considering relevant confounders revealed no significant diagnosis effect or interaction between diagnosis and region pair. The region pair of CA1 and SUB had significantly higher transcriptomic coupling than the CA1 with DLPFC (*F*(2)=7.714, *p* < 0.01, SUB_CA1-CA1_DLPFC Tukey p <.001), consistent with the monosynaptic nature of this circuit and higher similarity in cytoarchitecture between these two regions.

**Figure 4.**
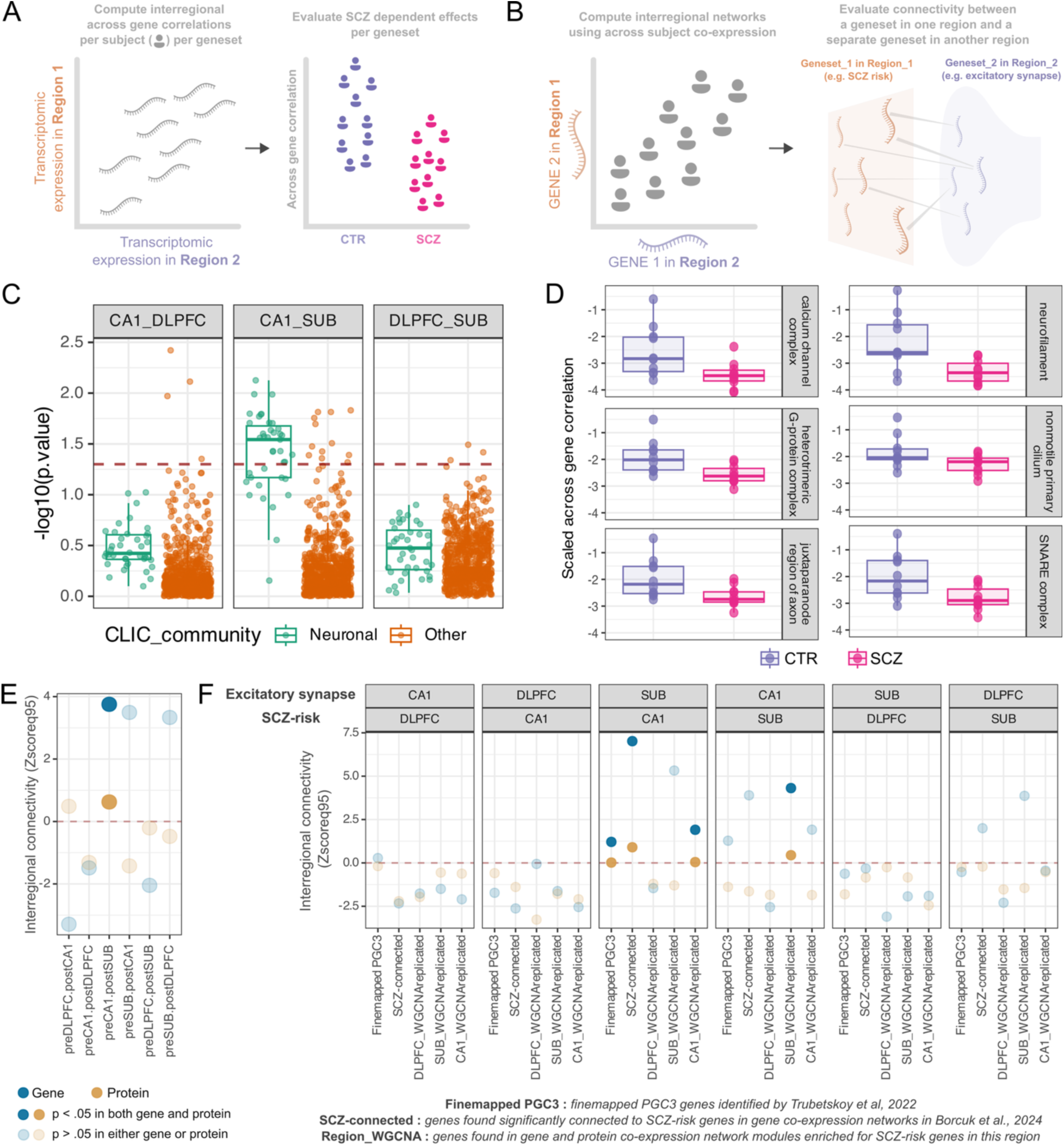
Interregional circuit-level effects of SCZ diagnosis and risk. **A)** To evaluate subject-level interregional transcriptomic coupling, we computed an across-gene correlation strength (Pearson R) per subject per region pair per gene set. Diagnosis effects of transcriptomic coupling were then assessed by comparing SCZ and CTR subjects using a linear model (relevant results shown in panels C and D). **B)** To evaluate interregional effects of SCZ risk on excitatory synaptic function, we constructed between-region across-subject co-expression networks per region pair and evaluated connectivity between one gene set (e.g., SCZ risk genes) in one region and a separate gene set (e.g., excitatory synaptic genes) in another region (relevant results shown in panels E and F). **C)** The diagnosis effect-log10(p-value) of linear models comparing transcriptomic coupling between SCZ and CTR subjects. The dashed red line reflects nominal significance. Points represent gene sets jittered along the x-axis; neuronal gene sets are colored in green, and all other gene sets are colored in orange. **D)** The six neuronal gene sets with the strongest diagnosis effects between SUB and CA1 are shown here, each showing decreased transcriptomic coupling between the SUB and CA1 in patients with SCZ at nominal significance. Points are subjects, colored, and spread on the x-axis by diagnosis. The y-axis indicates the across-gene correlation of a gene set scaled to all other across-gene correlation values within-subject. **E, F)** Interregional connectivity results between differing gene sets. Zscoreq95 indicates how many standard deviations the real connectivity value is above the 95^th^ quantile of the permuted null distribution, a value above 0 (red dashed line) representing an empirical one-sided p <.05. Blue points indicate results derived from gene networks, and tan points from protein networks. Dark points indicate where results are significant, p <.05 in both gene and protein networks. **E)** Interregional connectivity between pre-synaptic genes (*pre* prefix in x label, “presynaptic membrane” CLIC set) and post-synaptic genes (*post* prefix in x label, “dendrite membrane” CLIC set). In the x-axis, labels indicate which gene set is being evaluated in which region (e.g., *preCA.postSUB* indicates connectivity of pre-synaptic genes in CA1 with post-synaptic genes in SUB). **F)** Interregional connectivity between SCZ risk gene sets (*scz* prefix in bottom facet title) and excitatory synaptic genes (*exc* prefix in facet title, “excitatory synapse” CLIC set). The top two facet titles simply highlight which regions are being evaluated. The bottom facet titles indicate which gene set is being evaluated in which region (e.g., *excSUB.sczCA1* indicates connectivity of excitatory synaptic genes in SUB with SCZ risk genes in CA1). The x labels indicate the specific SCZ risk set being evaluated (*Finemapped PGC*: finemapped PGC3 genes identified by Trubetskoy et al, 2022 ^1^, *SCZ-connected*: genes consistently most connected to prioritized PGC3 genes in DLPFC and or HP networks identified by Borcuk et al., 2024 ^50^, *REGION_WGCNAreplicated*: union of genes found in WGCNA SCZ risk modules across both transcript and protein networks described in Figure 3D). Abbreviations: CLIC, CLustering by Inferred Co-expression; SCZ, schizophrenia; CTR, control.

We then considered transcriptomic coupling at the level of specific subsets of genes. Neuronal gene sets showed lower across-gene correlation strength than all other CLIC sets, on average (Figure S4B), possibly in association with the interregional variability of neurons compared to glia. We interrogated whether certain transcriptomic gene sets, such as neuronal sets, showed diagnosis effects by comparing within-subject scaled across-gene correlation values between SCZ patients and CTR subjects for each gene set, using a linear model to account for subject-wise confounders. Neuronal gene sets in general were descriptively more strongly correlated in CTR subjects between CA1 and SUB than in patients (Figure 4C, median nominal p value for neuronal gene sets < 0.05). Figure 4D shows the top six strongest differentially correlated neuronal gene sets arising from the SUB-CA1 connection (all p <.05, all t(20) > 2.6). Our results hint towards a reduced intra-hippocampal similarity and/or coordination of neuronal gene sets in SCZ patients, but are not significant after multiple comparison correction, likely due to the small number of individuals (n=20) and are therefore considered exploratory.

### Interregional circuit-level effects of SCZ risk on excitatory synaptic function

Considering that these data arise from glutamatergic neuron-enriched samples, we were interested in how SCZ risk in one region may affect excitatory synaptic function in another. We constructed between-region across-subject co-expression networks and evaluated interregional connectivity between SCZ risk genes and excitatory synaptic genes (Figure 4B). Interregional co-expression networks were constructed for each region pair using gene and protein data, keeping only genes/proteins that were present across both domains and in all three regions. Networks were thresholded to keep only the top 10% of connections. For each region-pair, we computed the number of retained connections between SCZ risk genes in one region and excitatory synaptic genes of another region. This value was compared to a distribution of connectivity to 1000 null risk sets of the same size and degree distribution as the original set. The Zscoreq95 value describes how many standard deviations the real value is above the 95^th^ quantile of the null. Values above 0 in this measure represent an empirical one-sided nominal p <.05. We prioritized results showing significance in both gene and protein-derived networks (dark points in Figure 4E, F).

To assess the validity of this approach in capturing biologically coherent signals, we first performed a preliminary evaluation of how pre-synaptic genes (“presynaptic membrane” CLIC set) connect to post-synaptic genes (“dendrite membrane” CLIC set) interregionally. The only significant result in both protein and gene networks was between CA1 pre-synaptic genes and SUB post-synaptic genes (Figure 4E, dark points), a coherent finding likely driven by the prominence of monosynaptic CA1->SUB excitatory efferent projections in the hippocampal circuit ^49^.

We then evaluated the connectivity of SCZ risk sets in one region with “excitatory synapse” CLIC genes of another region. We evaluated various types of SCZ risk sets. As a proxy of pure SCZ risk, we used the set of all fine-mapped PGC3 genes ^1^(*Finemapped PGC3*). Additionally, to proxy a stronger co-expression risk signal, we used a recently published set of genes that are significantly connected to PGC3 genes in DLPFC and/or HP tissue across various consortia ^50^ (*SCZ-connected*). Finally, for each region, we used the union of all genes in our SCZ risk modules found replicable across transcript and protein WGCNA networks (*REGION_WGCNAreplicated*, the per-region union of those genes described in Figure 3D). From the CA1, we found significant interregional connectivity of several SCZ risk gene sets (*Finemapped PGC3, SCZ-connected,* and *CA1_WGCNAreplicated)* to excitatory synaptic genes in the SUB (Figure 4F, third panel). From the SUB, only the *SUB_WGCNAreplicated* gene set, enriched for post-synaptic genes (see Figure 3D), was strongly connected to excitatory synaptic genes in the CA1(Figure 4F, fourth panel). In summary, we show that SCZ risk converges on directionally specific excitatory synaptic processes within the hippocampal formation (here CA1 to SUB).

### SCZ-dependent changes in cell type interactions within the CA1

We have so far highlighted an especially strong role of SCZ risk in CA1 and have detailed how this may affect CA1 excitatory neurons and their downstream synaptic properties at SUB. Given this result, we further assessed how interactions between excitatory neurons and other cell-types in CA1 are affected in SCZ. snRNA-seq was performed on CA1 tissue, and cell type clusters were identified using previously defined markers from hippocampal snRNA-seq ^20^ (Figure 5A). We evaluated enrichment of SCZ risk gene sets in each cell type cluster, using AUCell, which calculates enrichment as the proportional area under the curve of gene expression rankings (AUC) (Figure 5B). We assessed the same SCZ risk gene sets as described in the preceding sections, except we used a set of genes significantly connected to PGC3 genes specifically in HP tissue (*SCZ-connected_HP*). Gene sets consisting of SCZ-partners derived from co-expression (*SCZ-connected_HP, REGION_WGCNAreplicated*) showed stronger AUC enrichment values in neurons, more strongly excitatory neurons, as compared to the pure SCZ risk gene set (*Finemapped PGC3*). Moreover, the LCM-derived SCZ-partners (*REGION_WGCNAreplicated)* are also more strongly enriched in neuronal cell types than bulk-derived SCZ-partners (*SCZ-connected_HP)*. SCZ-dependent DEGs were found most prominently in excitatory cell types (Figure 5C), with both down (Figure 5D) and up (Figure 5E) regulated genes showing synaptic ontologies. We used *CellChat* ^51^ to identify cell type ligand-receptor pathway interactions that are differentially regulated in SCZ. In SCZ samples, there was an increased variety of significant interactions between glial and neuronal cell types and a decreased number of significant interactions between inhibitory and excitatory neuronal cell types as compared to CTR (Figure 5F). SCZ up-regulated DEGs present in SCZ-specific pathways are related to cell junction and adhesion. Instead, SCZ down-regulated DEGs present in CTR-specific pathways were related to GABA-ergic synapses (Figure 5G). We more specifically interrogated canonical inhibitory neuropeptide inputs to Excit_A, the cell type cluster found significantly enriched in CA1 LCM tissue (see Figure 1D), and found an absence of significant somatostatin and neuropeptide Y (NPY) pathways in SCZ patients (Figure 5H). Altogether, these results show increased neuronal expression of SCZ risk co-expression partners, increased involvement of glial cells, and decreased involvement of inhibitory cells with excitatory clusters in SCZ in the CA1.

**Figure 5.**
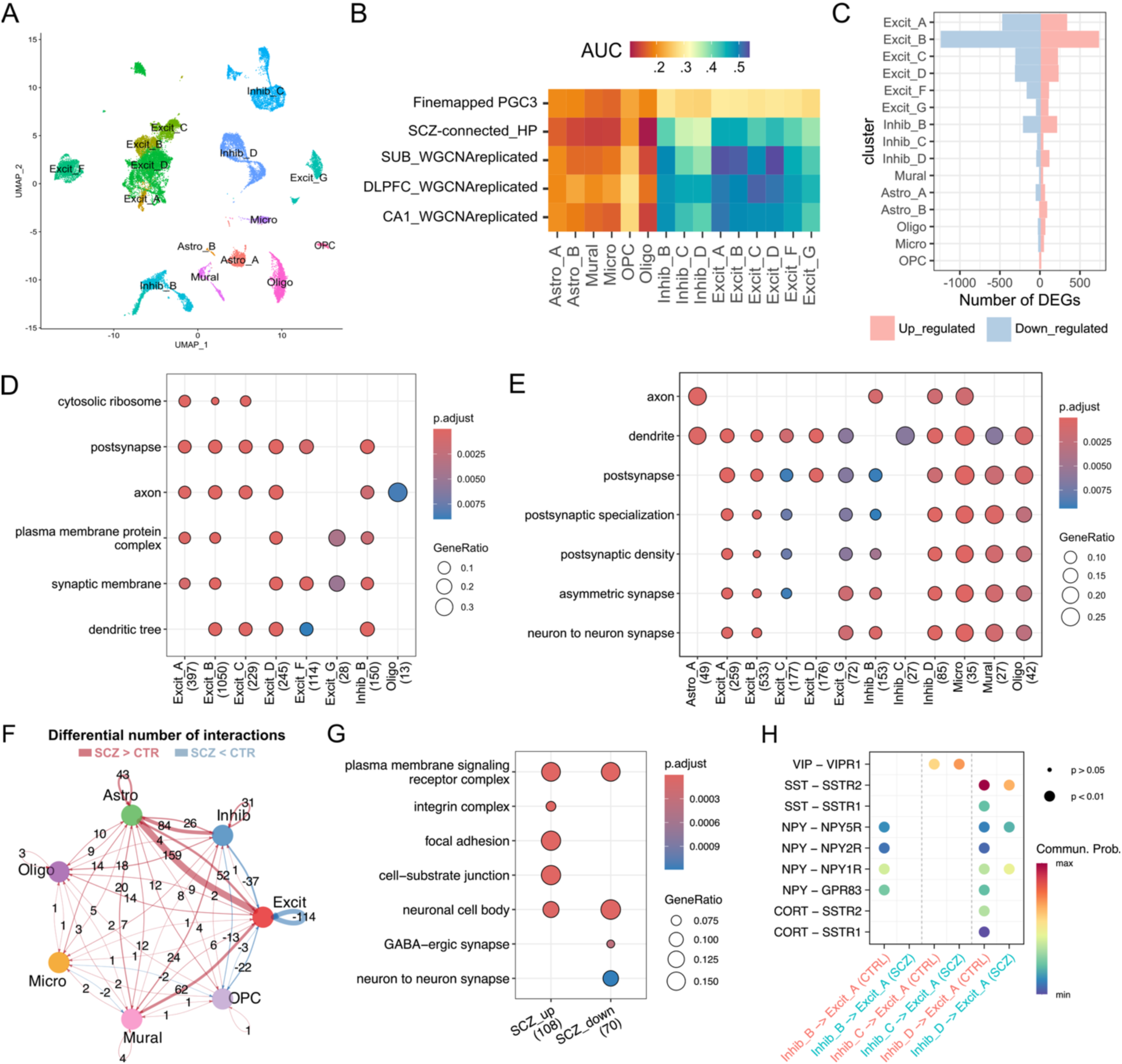
SCZ-dependent changes in cell type interactions within the CA1. **A)** We performed snRNA-seq on the CA1 in CTR subjects and SCZ patients. **B)** AUCell enrichment was used to evaluate the enrichment of SCZ risk gene sets (y-axis) in cell type clusters (x-axis). The color indicated the AUC value, which represents the proportional area under the curve of gene expression rankings for a specific gene set in a specific cell type cluster. **C)** SCZ-dependent DEGs were identified per cell type. **D, E)** GO enrichments for down **(D)** and up-regulated **(E)** genes are strongly synaptic regardless of cell type. **F, G, H)** We used *CellChat* to determine if certain cell types show differential ligand-receptor pair interactions in SCZ. **F)** In SCZ subjects, we find more ligand receptor interactions (red lines: SCZ > CTR) between glial and neuronal clusters and fewer interactions between inhibitory and excitatory neuron clusters (blue lines: SCZ < CTR) as compared to CTR subjects. **G)** The ligands/receptors in SCZ upregulated interactions are again enriched for terms related to cell junction and adhesion, while SCZ downregulated interactions are enriched for synapse, specifically GABA-ergic synapse. **H)** We took a closer look at which classical GABA-ergic neuropeptide inputs from inhibitory clusters to the Excit_A cluster (the cluster that shows a significantly enriched proportion in the LCM sample, see Figure 1D) were altered in patients and found absent or reduced NPY and somatostatin (SST) signaling. Abbreviations: DEG, Differentially expressed gene; SCZ, schizophrenia; CTR, control; NPY, neuropeptide Y; SST, somatostatin; VIP, Vasoactive intestinal peptide; CORT, coristatin; OPC, oligodendrocyte progenitor cell.

## Discussion

We used LCM to collect excitatory neuron-enriched samples from mono-synaptically linked nodes of a SCZ-associated neural circuit, i.e., from glutamatergic neurons in the CA1 hippocampal field, in SUB, and in layer III of DLPFC. LCM and bulk tissue expression were compared using a variety of quality control analyses. As expected, LCM showed increased expression of genes related to synaptic ontologies, and an enrichment of coherent excitatory neuronal sub-types based on reference DLPFC and HP snRNA-seq and DLPFC spatial transcriptomic data. A machine learning approach showed that LCM expression predicts regional identity better than bulk tissue expression, while protein and transcript expression predict regional identity better than gene expression. The translatability of a gene from transcriptomic to proteomic states was shown to at least partly depend on the stability of that gene’s transcriptomic expression over time, as assessed in iPSC-derived neurons. Effects of diagnosis on gene stability were examined, revealing relatively increased stability of cell junction processes and decreased stability of synaptic processes in iPSC derived neurons from SCZ subjects. We also examined diagnosis effects on protein phosphorylation, showing that phosphorylated peptides are generally increased in SCZ patients, specifically in the hippocampal subregions. In each region, co-expression networks were then constructed, and co-expression partners of SCZ risk genes were prioritized as those replicated across transcript and protein assays. Replicated SCZ risk partners showed consistent ontologies related to synaptic function, SUB more specific to post-synaptic glutamatergic function, and in the DLPFC and CA1, more related to transmembrane transporter activity.

We further evaluated interregional circuit-level effects of SCZ diagnosis and risk, using exploratory analyses that leverage our regionally overlapping expression data. The CA1-SUB circuit was particularly affected, with decreased interregional connectivity of neuronal gene sets in SCZ patients as compared to CTR subjects, and significantly decreased connectivity of SCZ risk genes in the CA1 with excitatory synaptic genes in the SUB. Finally, we used snRNA-seq data to examine the effect of SCZ diagnosis on cell type interactions within the CA1, where CA1 excitatory neurons showed an increased number of interactions with glial cells and a decreased number of interactions with inhibitory neurons in samples from patients with SCZ.

Though LCM generally demonstrated greater proportions of coherent neuronal excitatory subtypes compared with bulk tissue, there appeared to be a certain ambiguity in the genetic signature of L3 DLPFC neurons, mapping to an L4 cluster rather than L3. The ambiguity of L3 is not surprising, as L3 is sometimes grouped with L2 in DLPFC snRNA-seq data ^34,52^. This ambiguity was also noted by Maynard et al., who found that layer 3 had the fewest specific genes compared to other layers in their spatial transcriptomic study ^3^. Projection targets of a DLPFC subfield greatly influence its transcriptome ^53^. Since circuit structure is more clearly defined in the HP (e.g., prominence of mossy fibers and Schaffer collaterals), snRNA-seq clusters of the DLPFC may be more challenging to resolve via computational deconvolution than clusters of hippocampal cells.

We prioritized consensus SCZ risk partner genes found in SCZ risk-enriched co-expression modules that are present across both the transcriptome and the proteome. Solute carrier proteins (SLC) made up five of the eight genes found as consensus SCZ risk partners in at least two regions, always in CA1. This family of proteins is responsible for the transmembrane movement of nutrients and ions, and plays an important role in maintaining cell homeostasis ^42^. SLC genes have previously been linked to SCZ-dependent effects in the HP, showing SCZ-dependent upregulation in the CA1 that is rescued by anti-psychotic treatment ^54^. These results together suggest that disrupted homeostasis of the HP via dysregulation of SLC genes potentially contributes to SCZ.

An interesting entry among these genes was *SLC12A5,* found to be a SCZ partner gene in both DLPFC and CA1. This gene encodes the K+-Cl− cotransporter 2 (KCC2), the main Cl^-^ exporter in neurons ^55^. *SLC12A5* has been found significantly connected in coexpression networks to PGC3 genes in the DLPFC and HP ^50^. Expression of *SLC12A5* was found to be affected in SCZ patients, with decreased expression of the full-length *SLC12A5* in the HP ^56^, and increased expression of the *EXON6B* isoform in the DLPFC ^57^. *SLC12A5* is linked to the regulation of inhibitory function and is a principal driver of switching excitatory-to-inhibitory GABA polarity through neurodevelopment, a phenomenon that has been reported to be altered in association with SCZ ^58^. In a similar vein, we also identified *SLC4A3*, which encodes the anion exchange protein 3 (AE3), a Cl−/HCO3− exchanger also shown to affect GABA function ^59,60^.

*SLC12A5* also plays a direct role in post-synaptic glutamatergic function, where it has been shown to regulate the formation of glutamatergic synapses through cytoskeletal coupling in dendritic spines ^61^. *SLC6A17,* identified in the CA1 and DLPFC, is involved in the presynaptic vesicular reuptake of amino acids such as glycine and glutamate, and is also shown to regulate glutamatergic synapse formation ^62,63^. A more direct link to glutamatergic synapse dysfunction was identification of *GRIA2* as a SCZ risk partner in DLPFC and SUB subfields, which encodes the GluR-2 subunit of the AMPA receptor. This finding resonates with a recent review that highlighted consistent decreased expression of *GRIA2* in SCZ across 4/6 published studies of medial temporal lobe regions ^43^. Another link to AMPA receptor dysfunction is *LRRTM2*, identified in the CA1 and SUB, which is essential for the positioning of AMPA receptors in the postsynapse of hippocampal neurons ^44–46^. In summary, our SCZ risk partners highlight potential mechanisms by which excitatory synapses may be affected in SCZ, such as by dysregulation of AMPA receptors via *GRIA2*/*LRRTM2* or altered regulation of glutamatergic synapse formation by *SLC12A5/SLC6A17*.

We continued this line of investigation, interrogating how SCZ risk further affects excitatory synaptic function cross-regionally. Using three different SCZ risk gene sets, CA1 SCZ risk had consistently significant connectivity to SUB excitatory synaptic genes in both gene and protein-derived interregional networks, highlighting that SCZ risk in this region may have a higher propensity of affecting downstream rather than upstream regions of the circuit. Two of these SCZ risk gene sets contain *SLC12A5*, the *SCZ-connected* and *CA1_WGCNAreplicated* sets, hinting that CA1-SUB glutamatergic projections may be strongly affected by *SLC12A5* dysregulation in CA1. We also found that the *SUB_WGCNAreplicated* genes in SUB, containing mainly post-synaptic genes (see Figure 3d), including *GRIA2* and *LRRTM2,* are strongly connected to CA1 excitatory synaptic genes. This, in turn, suggests that post-synaptic and AMPA receptor-related dysfunction in the SUB may work in concert with CA1 synaptic expression to drive the CA1->SUB circuit dysfunction.

Interregional co-expression of gene expression is shown to associate with interregional functional connectivity ^48^, and our analysis may help point to potential mechanisms driving dysfunctional connectivity of the HP in SCZ ^47,64^. Although disrupted functional connectivity between the DLPFC and the HP is a consistent phenotype in SCZ ^25,31^, we did not find any gene-protein replicable effects of SCZ risk-related interregional connectivity between the DLPFC and HP, with only gene-level effects identified between SCZ risk in SUB and excitatory synapse in DLPFC. The protein-derived effect of SCZ risk on excitatory synaptic function appears to be generally weaker than the gene-derived. This may be explained by a more dispersed localization of proteins, given that synaptic protein translation may occur locally in the neurite/synapse rather than in the nucleus or cell body ^65^. Regardless, significant protein-derived connectivity between SCZ risk and excitatory synapse was always accompanied by a stronger gene-derived effect, suggesting that protein-derived effects likely represent weaker representations of the same gene effects, at least as captured with our technique.

Our results overall show that SCZ risk processes in CA1 excitatory neurons may have implications for SUB synaptic processes downstream. snRNA-seq of the CA1 was utilized to further investigate how other cell types may affect excitatory neurons within CA1 of SCZ patients. SCZ patients showed increased interactions of excitatory neurons with glial cells in pathways related to the cell junction. This result replicates the findings of a more extensive study of DLPFC snRNA-seq^52^, which found a SCZ-dependent increase of glial-neuronal interactions using the same *CellChat* paradigm. This also corroborates our results showing an increased stability of cell junction terms in iPSCs derived from SCZ patients. However, these results contrast with a previous study that showed decreased expression of a neuron-glia concerted co-expression pathway in SCZ patients ^66^, suggesting that at least in DLPFC ligand-receptor interactions and concerted co-expression may represent different processes altered in SCZ (e.g., increased astrocytic adhesion but reduced synapse formation). Our snRNAseq data also revealed a SCZ-associated decrease in interactions between excitatory neurons and inhibitory neurons. This may help drive disrupted excitatory inhibitory balance, a common phenotype of SCZ ^67^ and a potential mechanism by which connectivity with the CA1 is affected ^68,69^. *SLC12A5* or *SLC4A3* may also play a role in disrupting GABAergic function here ^55,59,60^, an example of how identified SCZ risk partners may affect SCZ phenotypes in this snRNA-seq data.

Our study comes with limitations. The small sample size limited our ability to generate per-gene or per-module diagnosis-driven statistics, such as DEGs, or detecting diagnosis-associated module eigengenes as is typically done in WGCNA studies, and to test for interactions of diagnosis and regional circuitry. Regardless, preliminary results are encouraging for further investigations of interconnected circuits involved in SCZ. As LCM is not a ‘single cell’ technique at present, but depends on pools of similar cells identified either by morphology or specific cell markers, caution is warranted when interpreting our results as reflecting a glutamatergic-only neuronal circuit. Another limitation is the use of short-read sequencing and computational isoform transcript reconstruction. Yet, our results show that transcript reconstruction still increases information content for regional identity discrimination.

In summary, our study highlights high-fidelity gene-to-protein replicable mechanisms of SCZ risk in a circuit linking excitatory neurons of hippocampal subregions and DLPFC. We demonstrate a possible flow of these SCZ risk pathways to affect downstream synaptic processes, specifically from CA1 efferents to tSUB. We show how these may be related to common phenotypes associated with SCZ, such as hippocampal disruptions in functional connectivity, or disrupted excitatory-inhibitory balance.

## Materials and Methods

For information on subjects and tissue samples used, see Table 1.

**Table 1.** Human brain subject demographics. SCZ=schizophrenia. CTR=neurotypical control. L=Left. R=Right. RIN= RNA Integrity Number, PMI= postmortem interval. All individuals were of male sex. Brain regions included per subject: dorsolateral prefrontal cortex, cornu ammonis of hippocampal formation pyramidal CA1 neurons, large pyramidal neuronal layer of presubiculum. *Sampled from the right hemisphere. **History of smoking at time of death reported by next of kin.

| Diagnosis | Age at death (years) | DLPFC RIN | PMI (hours) | Manner of Death | History of Smoking** | Nicotine/ Cotinine by (Toxicology) | Antipsychotics at time of death (Toxicology) | Comorbid Substance Use Disorder (type) |
| --- | --- | --- | --- | --- | --- | --- | --- | --- |
| SCZ | 30.0 | 8.5 | 23.0 | Suicide | No | Negative | Yes (None) | No |
| SCZ | 46.1 | 6.7 | 22.5 | Natural | No | Negative | Clozapine | No |
| SCZ | 65.3 | 8.8 | 15.5 | Natural | Yes | Positive | Chlorpromazine | No |
| SCZ | 51.2 | 6.7 | 23.0 | Undetermined | Yes | Positive | Olanzapine | Alcohol |
| SCZ | 26.2 | 7.7 | 28.5 | Suicide | Yes | Positive | Haloperidol, Olanzapine | Cannabis, Opioid |
| SCZ | 67.0 | 7.2 | 36.5 | Natural | Yes | Positive | Risperidone | Alcohol |
| SCZ | 23.1 | 8.3 | 32.5 | Suicide | No | Negative | Yes (None) | Cannabis, Hallucinogen |
| SCZ | 60.7 | 7.2 | 28.0 | Natural | No | Negative | Lurasidone | Alcohol, Sedative |
| SCZ* | 49.1 | 6.9 | 41.0 | Natural | No | Negative | Risperidone, Paliperidone | No |
| SCZ | 60.1 | 7.3 | 38.5 | Natural | No | Negative | Yes (None) | No |
| CTR | 49.3 | 8.4 | 29.5 | Accident | No | Negative | Negative | No |
| CTR | 42.1 | 7.5 | 28.5 | Natural | No | Negative | Negative | No |
| CTR | 62.0 | 6.5 | 29.0 | Natural | No | Negative | Negative | No |
| CTR | 51.6 | 8.5 | 38.5 | Accident | Yes | Positive | Negative | No |
| CTR | 47.8 | 7.1 | 25.5 | Natural | No | Negative | Negative | No |
| CTR | 72.4 | 7.4 | 27.5 | Accident | Yes | Negative | Negative | No |
| CTR | 61.8 | 7 | 22.5 | Natural | Yes | Positive | Negative | No |
| CTR | 59.9 | 7.4 | 26.0 | Natural | No | Negative | Negative | No |
| CTR | 34.6 | 8.3 | 36.0 | Accident | No | Negative | Negative | No |
| CTR | 52.7 | 8 | 28.0 | Accident | No | Negative | Negative | No |

### Human postmortem brain tissue acquisition and dissection

The research described herein complies with all relevant ethical regulations. Postmortem human brain tissue was obtained as previously described (Collado-Torres et al., 2019). Briefly, tissue was collected at several sites for this study. Two human brain subjects were obtained at the Clinical Brain Disorders Branch at the NIMH from the Northern Virginia and District of Columbia Medical Examiners’ Office, in accordance with NIH Institutional Review Board guidelines (protocol 90-M-0142). These samples were transferred to the LIBD under a material transfer agreement with the NIMH. The remaining samples were collected at the LIBD according to a protocol approved by the Institutional Review Board of the State of Maryland Department of Health and Mental Hygiene (12-24) and the Western Institutional Review Board (20111080).

Audiotaped informed consent to study brain tissue was obtained from the legal next of kin on every case collected at the NIMH and LIBD. Details of the donation process and specimen handling have been described previously (Lipska BK et al., 2006). After obtaininginformed consent to brain donation, a standardized 36-item telephone screening interview was conducted (the LIBD autopsy questionnaire) to gather additional demographic data, along with detailed psychiatric, neurological, substance use, treatment, general medical, family and social histories. A clinical narrative summary was written for every donor to include data from multiple sources, including the autopsy questionnaire, medical examiner documents (investigative reports, autopsy reports and toxicology testing), macroscopic and microscopic neuropathological examinations of the brain and extensive psychiatric, neurologic, and medical record reviews, supplemented with additional family informant interviews using the mini-international neuropsychiatric interview. Two board-certified psychiatrists independently reviewed every clinical case summary to arrive at DSM-V lifetime psychiatric, and substance use disorder diagnoses, including SCZ and bipolar disorder, as well as substance use disorders, and if for any reason agreement was not reached between the two reviewers, a third board-certified psychiatrist was consulted.

All donors were free from significant neuropathology, including cerebrovascular accidents and neurodegenerative diseases. Available postmortem samples were selected based on RNA quality (RIN ≥ 6.5).

A toxicological analysis was performed in each case. The non-psychiatric non-neurological neurotypical individuals had no known history of significant psychiatric or neurological illnesses, including substance abuse. Positive toxicology was exclusionary for neurotypical individuals but not for individuals with psychiatric disorders.

### Human postmortem brain processing, dissections, and donor subject details

Postmortem fresh human brain dissections and freezing were performed as described previously (Collado-Torres et al., 2019). Anatomically, two frozen blocks containing – (i) the dorsolateral prefrontal cortex (Brodmann areas 9/46) and (ii) the presubiculum and cornu ammonis CA1 of the human hippocampal formation at the level of the lateral geniculate nucleus – were dissected from each of the 20 subject brains. The blocks were dissected out from previously fresh frozen coronal 0.75 cm thick human brain slices using a hand-held dental drill (Cat# UP500-UG33, Brasseler, Savannah, Georgia) as described before (Lipska BK et al., 2006). Specific gross anatomical landmarks above were matched to corresponding coronal brain sections from the Allen Human Brain Reference Atlas (https://atlas.brain-map.org) for each subject to ensure anatomic consistency along the rostrocaudal axis of each brain region.

The demographic data are detailed by each subject in Table 2 Table 2 includes individual-level demographic information, including sex, ancestry, age, postmortem interval, manner of death and treatment history for each of the donors.

**Table 2:**
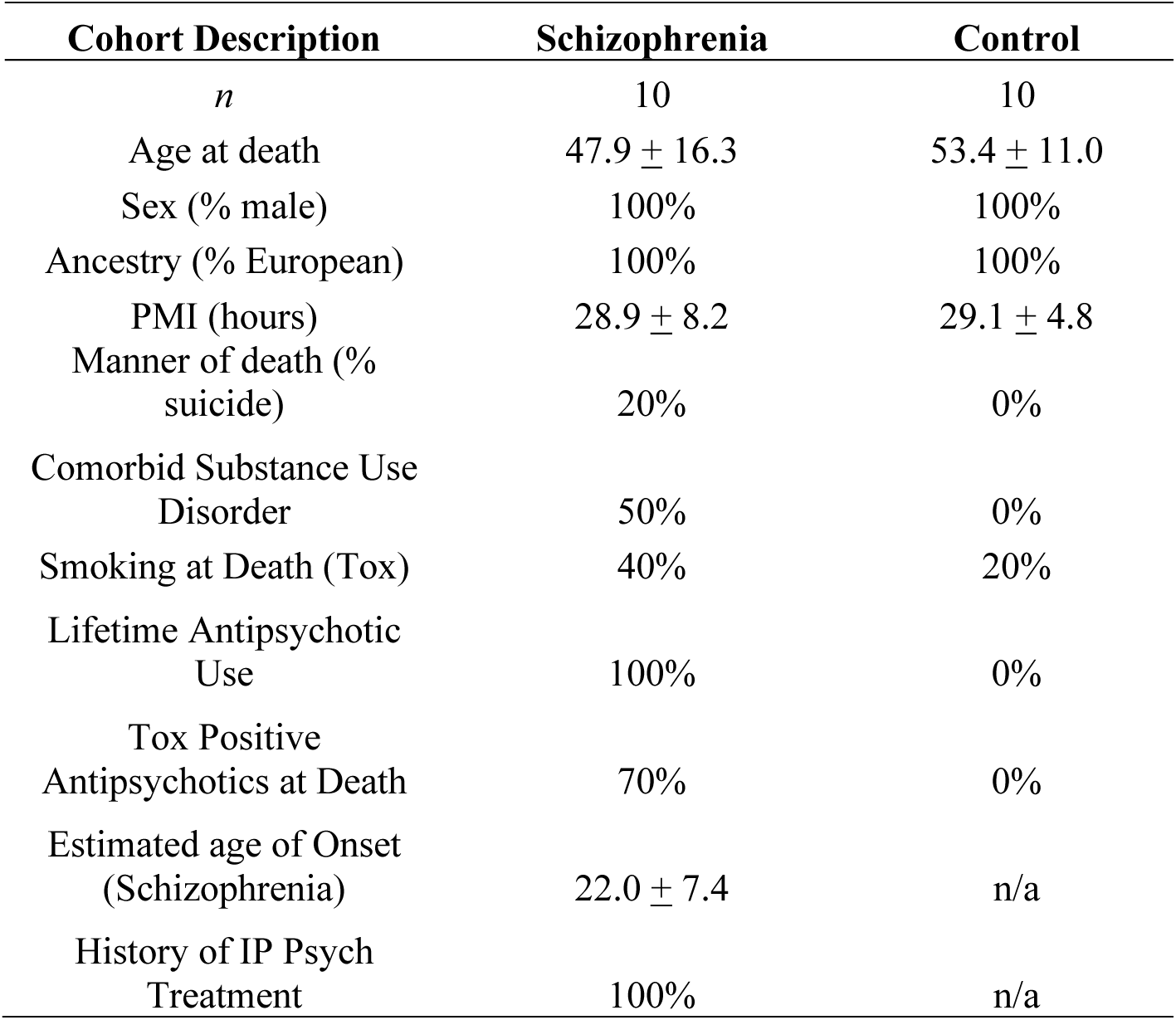
Summary Table of human brain subjects.

| Cohort Description | Schizophrenia | Control |
| --- | --- | --- |
| *n* | 10 | 10 |
| Age at death | 47.9 ± 16.3 | 53.4 ± 11.0 |
| Sex (% male) | 100% | 100% |
| Ancestry (% European) | 100% | 100% |
| PMI (hours) | 28.9 ± 8.2 | 29.1 ± 4.8 |
| Manner of death (% suicide) | 20% | 0% |
| Comorbid Substance Use Disorder | 50% | 0% |
| Smoking at Death (Tox) | 40% | 20% |
| Lifetime Antipsychotic Use | 100% | 0% |
| Tox Positive Antipsychotics at Death | 70% | 0% |
| Estimated age of Onset (Schizophrenia) | 22.0 ± 7.4 | n/a |
| History of IP Psych Treatment | 100% | n/a |

### Laser capture microdissection

All brain sections were collected using the Leica 350 cryostat precooled to-20C at the level of the lateral geniculate nucleus of the thalamus for the hippocampal formation (including dentate gyrus, CA3/2/1, presubiculum, and subiculum) and BA9/46 DLPFC for every subject (Table 1). Every section was obtained at a thickness ranging between 25-40 microns and transferred onto glass slides coated with precharged PEN membrane (Zeiss # 415190-9081-000). An average of 10 sections corresponding to ∼5000 layer 3 pyramidal neuron enrichments, ∼5000-7000 pyramidal neurons of the CA1, and ∼5000-7000 pre-subiculum large pyramidal neuron enrichments were harvested from every subject. All sections were uniformly Nissl stained using the ThermoFisher Arcturus Histogene staining kit (KIT0401). Nissl stained sections were compared to anatomical Nissl sections of the human Allen Brain Atlas for dlpfc (BA9/46), CA1 stratum pyramidale, and subicular formation for the major pyramidal layer (https://atlas.brain-map.org) on every subject to ensure anatomical consistency before neuron enrichments were cut using laser capture microdissection as described previously (Jaffe AE, Hoeppner DJ et al., 2020). Nissl-stained sections were used to determine the boundaries of layer 3 pyramidal neurons. Starting from the apical surface of the neocortex (layer 1), layer 3 pyramidal neurons were distinctly identified by lower-density larger neuronal cell bodies relative to both layers 2 and 4. This approach has been successfully implemented with the Leica laser capture system to cut out the entire layer 3 cells (MacDonald ML et al., 2019). Here, the Zeiss PALM system additionally offers the feature to selectively highlight neuronal cells within neocortical layers for microdissection, which facilitated selective enrichment for layer 3 pyramidal neurons. Specific pyramidal neurons were highlighted manually and entered into the Zeiss PALM Robo v4.9 LCM system. The Zeiss cool UV laser (355nm) uses photon excitation energy to catapult selected neurons onto Zeiss LCM tube caps (Carl Zeiss™ 415190-9211-000) and therefore never comes in direct contact with the tissue. The no-contact high accuracy Zeiss PALM microbeam features allows for selective enrichment of morphologically defined neurons with corresponding high extraction yields for both RNA and protein from the same neuron lysate. Once the neurons were highlighted, cut, and catapulted onto LCM adhesive caps, the caps were immediately stored on dry ice until lysis and RNA extraction by RNeasy Micro kit (Qiagen). Over 250 ng of total RNA was isolated from the pooled neuron enrichments for each donor. This relatively high output quantity from LCM enables more accurate steady-state mRNA quantification and avoids the amplification bias and computational confounders associated with RNA preamplification (Ziegenhain C et al., 2017). The quantity and integrity of the RNA were determined by NanoDrop and BioAnalyzer (Agilent).

### RNA sequencing

Briefly, 200 ng RNA from every sample was utilized for RNA-sequencing without any pre-amplification as described previously ^2^. Sequencing libraries were prepared using the TruSeq Stranded Total RNA Library Preparation kit with RiboZero Gold rRNA depletion. For quality control, synthetic External RNA Controls Consortium (ERCC) RNA Mix 1 was spiked into each sample. Samples were balanced for diagnosis within each batch. The resulting paired-end, strand-specific libraries were sequenced on an Illumina HiSeq3000 at the LIBD Genomics Sequencing Facility. FASTQ files were generated using the Illumina Real-Time Analysis module by performing image analysis, base calling, and the BCL Converter (CASAVA v1.8.2). Alignment of reads was performed to the hg38/GRCh38 human genome (Gencode release 25, GRCh38.p7, chromosome only) using HISAT2 (v2.04) and Salmon (v0.7.2) using the reference transcriptome to initially guide alignment based on annotated transcripts. The synthetic ERCC transcripts were quantified with Kallisto (v0.43.0).

### RNA Data processing

Counts were generated as described previously. The sorted BAM files from HISAT2 alignments were generated and indexed using SAMtools (v1.6; HTSlibv1.6). The quality of alignment was assessed using RSeQC (v2.6.4). Transcriptomes were characterized for genes and transcripts.

Gene counts were generated using the SubRead utility featureCounts (v1.5.0-p3) for paired-end, reverse-stranded read counting. Transcript counts and transcripts per million (TPM) estimates were generated using Salmon. These estimated transcript count Salmon files were used for downstream differential expression analysis. Quality control of samples was determined based on ERCC concentrations, genome alignment rate, gene assignment rate (>20%), and mitochondrial mapping rate (<6%) as described previously (Jaffe AE, Hoeppner DJ et al., 2020).

### Direct Comparison of protein profiles by Tandem Mass Tag analysis

Sample preparation. A total of 60 neuronal lysates were processed for proteomics extractions from the organic phase (note that same neuronal lysate supernatant phase was used for RNA extractions and RNA sequencing as described above). Sample preparation. A total of 60 neuronal lysates were processed for proteomics extractions from the organic phase (note that same neuronal lysate supernatant phase were used for RNA extractions and RNA sequencing as described above). Isopropanol precipitated organic phase pellets were washed in absolute ethanol, then resuspended by pipetting in SDS (10%) and DTT (200mM) (Sigma, #43819) at 60C for 1 hour to reduce disulfide bonds. Lysate volumes were made up to 250μl Tri-ethyl ammonium bicarbonate buffer (100mM) (TEAB buffer, Sigma, #T7408) followed by gentle mixing, and incubation at-80C for 4 hours. Thawed lysates were then alkylated by iodoacetamide (250 mM) (Sigma #I6125) in the dark for 15 mins followed by centrifugation and protein quantification via BCA Assay (Pierce). Post-quantification, a total of 50 μg protein per sample was taken for tandem mass tag (TMT) labeling and dried by vacuum centrifugation.

Protein Digestion and TMT Labeling. All 60 protein samples were digested using a modified on-bead solid phase digestion protocol, as follows. Proteins were rehydrated with 10uL dithiothreitol (DTT, 50mM, Sigma, #43819) at 60C for 1 hour to reduce disulfide bonds, followed by alkylation with 10uL chloroacetamide (100mM, Sigma # **<u>22790</u>**). Proteins were then buffer exchanged using a modified solid phase extraction method, as described elsewhere ^70^. Each sample was diluted with 50ul ethanol, incubated with 10ul magnetic beads for 5 mins while gently shaking at 1,000rpm. Samples were placed onto a magnet and the supernatant was removed and discarded. The sample was washed with 180ul 80% ethanol, and steps were repeated three times. The proteins were then digested with 100ul of Trypsin/ LysC mixture (V5071, Promega, 20mg/2ml in 100mM TEAB) overnight at 37C while shaking. Post-digestion, individual samples were then labeled directly into the sample (TMT 10-plex, ThermoFisher Scientific) so every sample had a unique isobaric mass tag, as per manufacturer’s instructions. All samples within each of the 6 batches (10 samples per batch) were randomly paired and labeled. Each 10-plex batch had 5 control and 5 SCZ samples of similar brain regions to minimize variability and for better analysis design. Briefly, the TMT-10 plex reagents (0.8μg/vial) were brought to room temperature, and 41μl of anhydrous acetonitrile was added to each vial, vortexed, and centrifuged. An entire vial was added to each sample of 50ug tryptic peptide and left at room temperature for 1 hour. Next, 8μl of hydroxylamine (5%) was added to quench the reaction, followed by multiplex combining of batches (10 samples/batch) and dry vacuuming them by centrifugation. The combined TMT labeled peptides were resuspended in 1mL of 10mM TEAB and separated by basic reverse phase using an acetonitrile gradient (0-90% in 10mM TEAB on a 5μm C18 Waters XBridge column using Agilent 1200 capillary HPLC in normal flow mode and Agilent 1260 micro-fraction collector) into 84 fractions. Alternating rows were concatenated to produce 24 fractions. For the direct mass spectrometry runs, 50ul (aprox 4ug) was removed and speedvacced to dryness. The remainder of the sample was also dried and used for the phosphoproteomic sample prep. For enrichment, the 24 fraction were concatenated into 12 fractions during the resuspension with 100ul of enrichment solution (25mg/ml dihydroxybenzoic acid, 10mg/ml titanium dioxide in 80% acetonitrile, 3.5% trifloroacetic acid). Samples were vortexed at 1,000rpm for 2 hours. The incubated sample was added to the top of a house-made C18 stage tip (Empore C18, 3M), which acts a filter to collect the TiO2 phase and wash the peptides. The wash steps included three washes with 75ul of glycolic acid wash buffer (80%acetonitrile, 3.5% trifloroacetic acid, 1M glycolic acid), and one wash without the glycolic acid. Phosphopeptides were eluted using 50ul of elution buffer (fresh 30% acetonitrile, 3% NaOH) and repeated once. One additional wash with 50ul of 80%acetonitrile was also done to make sure no peptides remained bound to the C18 in the stage tip. The phosphopeptides were neutralized with 50ul of 10% formic acid and dried by vacuum centrifugation..

Tandem Mass Spectrometry –Fractions were resuspended in 20 μL loading buffer (2% acetonitrile in 0.1% formic acid) and analyzed by reverse phase liquid chromatography interfaced with tandem mass spectrometry (LC/MSMS) as described previously (Barber CN et al. 2022) using an Easy-LC 1200 HPLC system1 interfaced with an Orbitrap Fusion Lumos Tribrid Mass Spectrometer (ThermoFisher Scientific). Peptides (20% injected, aprox. 1ug on column) were loaded onto a C18 trap (S-10 μM, 120 Å, 75 μm × 2 cm, YMC, Japan) and subsequently separated on an in-house packed PicoFrit column (75 μm × 200 mm, 15 μ, +/−1 μm tip, New Objective) with C18 phase (ReproSil-Pur C18-AQ, 3 μm, 120 Å2) using 2–90% acetonitrile gradient at 300 nl/min over 120 min. Eluting peptides were sprayed at 2.0 kV directly into the Lumos. Survey scans (full ms) were acquired from 300 to 1700 m/z with data-dependent monitoring of up to 15 peptide masses (precursor ions), each individually isolated in a 1.2 Da window and fragmented using HCD activation collision energy 32 and 30 s dynamic exclusion. Precursor and the fragment ions were analyzed at resolutions 70 000 and 35 000, respectively, with automatic gain control (AGC) target values at 3e6 with 50 ms maximum injection time (IT) and 1e5 with 200 ms maximum IT, respectively.

Isotopically resolved masses in the precursor (MS) and fragmentation (MS/MS) spectra were analyzed with Proteome Discoverer (PD) software (v2.4, Thermo Scientific). All data were searched by Mascot (2.6.23) against the Refseq 2017_Complete database (57 479 sequences for taxonomy Homo sapiens) using criteria: sample’s species; trypsin digestion allowing one missed cleavage; N-terminal TMT label as fixed modifications; TMT label on lysine, methionine oxidation, asparagine, and glutamine deamidation, and cysteine carbamidomethylation as variable modifications. Peptide identifications were filtered at 1% false discovery rate (FDR) confidence threshold, based on a concatenated decoy database search, using the Proteome Discoverer. Proteome Discoverer uses only the peptide identifications with the highest Mascot score for the same peptide matched spectrum from the different extraction methods.

Protein quantification for analyses. Briefly, peptides identified with a 1% FDR confidence threshold as described above were considered for analysis. Reporter ions from MS/MS spectra with isolation interferences ≥ 30% were excluded from further analysis. Protein relative abundances were determined from the normalized log2 values of the above spectra. The reporter ion intensities were first transformed into log2 notation and for each sample the median value for each peptide was taken to represent that peptide. The values of all samples were then quantile normalized to minimize technical variation, such as differential amounts of material loading, prior to fold change analysis.

### Pre-processing

Gene and transcript-level mRNA expression was quantified as Transcripts per Million (TPMs) and annotated as total gene expression separately for each brain region, regardless of alternative transcript quantification using GENCODE release 25 (GRCh38.p7). We selected genes above median TPM of 0.1 and free of floor effects (maximum 0% of zeroes per gene); we log-transformed TPM values with an offset of 1, i.e., log2(TPM+1). After removing mitochondrial genes, datasets included a variable number of genes for different regions.

For proteins, raw abundance values were divided by the total sum of protein abundance across all genes, then multiplied by a million, giving counts per million (CPM). These values were log-transformed with an offset of 1 to give logCPM.

### CIBERSORTx deconvolution

SingleCellExperiment files containing snRNA-seq data for the HP and the DLPFC were taken from Tran et al., 2021 at https://github.com/LieberInstitute/10xPilot_snRNAseq-human, using the links under the “Processed data” header. For each region, the SingleCellExperiment was used to construct a matrix for cell type expression (nuclei per column, gene per row) with columns annotated for cell type, that was input into CIBERSORTx to create a signature matrix of cell type expression profiles, using default parameters (min expression =.75, replicates = 5, sampling =.5). The resulting signature matrix was to compute cell type proportions from bulk and LCM RPKM data (BULK DLPFC and HP ^6^, and LCM DG ^3^), using 100 permutations, without batch correction.

### Differential gene expression analysis and expression dispersion in LCM

To identify significantly differentially expressed genes between LCM and bulk tissue samples, we performed Differential Gene Expression (DGE) analysis by using linear mixed-effects modeling using the *voom* function from the *limma* package ^71^. To control for technical variations such as differences in library preparation methods, we normalized the data by applying correction factors to several key metrics. These included i) The rate at which reads mapped to the mitochondrial chromosome; ii) The proportion of reads assigned to rRNA (ribosomal RNA); iii) The overall percentage of reads that aligned to the reference genome and iv) The fraction of reads that mapped to exonic (protein-coding) regions. This normalization step helps to ensure that any biological differences we observe are not confounded by variations introduced during sample processing and sequencing. For each brain region, we compared the gene expression profiles between the corresponding LCM and bulk samples: Bulk DLPFC (n = 25587) vs LCM DLPFC (n = 22782), Bulk HP (n = 27079)vs LCM CA1 (n = 23416), Bulk HP (n = 27079) vs LCM SUB (n = 23454).

Genes with an adjusted p-value less than 0.05 were considered statistically significant and were selected for further analysis.

We designated genes as annotated to layer 3 based on a spatial transcriptomics analysis of the DLPFC by Maynard et al 2021, using their Supplementary Table 5 ^35^. Their approach identified genes that show significantly increased expression to one or more layers (e.g. L2 and L3) as compared to all other layers (e.g. L1, L4, L5, L6, white matter). We considered genes as annotated to layer 3 as long as there was a significant association to layer 3, even if the significant association includes other layers.

For each set of differential expression analysis (Bulk DLPFC (deg = 4336) vs LCM DLPFC (deg = 2607), Bulk HP (deg = 4295) vs LCM CA1 (deg = 2200), and Bulk HP (deg = 3512) vs LCM SUB (deg = 1774), the significant genes were split into two groups based on the sign of their log fold change (logFC) values. Genes with positive logFC values, indicate higher expression in the LCM samples compared to the bulk samples. Genes with negative logFC values, indicate lower expression in the LCM samples compared to the bulk samples. Then, we combined all significant genes from each set and performed GO enrichment analysis to identify over-represented cellular component terms in each group with background universe being all genes from each set. Resulting p-values from the GO enrichment analysis were adjusted for multiple testing using Benjamini-Hochberg correction to control the false discovery rate (FDR). We aimed to identify genes and cellular components that are differentially expressed between LCM and bulk samples in the DLPFC, CA1, and SUB regions.

### Machine Learning Prediction

We studied RNA-seq and TMT proteomics data obtained through LCM from post-mortem brain tissues of ten patients with SCZ and CTR subjects. As a comparison, we included RNA-seq data obtained through bulk tissue from 30 SCZ patients and 30 CTR, matched to the demographic characteristics of the LCM sample. After data preprocessing, we employed high-quality gene, transcript, and TMT proteomics reads derived from the DLPFC and hippocampal subregions, SUB and CA1, to assess regional identity and compare it to bulk tissue data from the DLPFC and the HP. We determined differentially expressed genes in a leave-one-out cross-validation framework and used a Random Forest classifier to predict the region of the left-out sample in 100 iterations to generate reliable distributions. We also compared prediction accuracies within the same samples between the different omics data. Finally, we determined gene-protein correlation within the most predictive transcripts and peptides across subjects.

### Gene to protein translatability

A gene-to-protein translatability was evaluated per gene as a robust linear model using gene logTPM values to predict protein logCPM values, including confounders of age, diagnosis, mitochondrial_gene_mapping rate (gene level), gene mapping rate, total_protein_count, and protein_sequencing_batch, using the rlm() function from the *MASS* package on R. The translatability score was considered as the t value associated with logCPM in this model.

### iPSC stability

The iPSC generation pipeline has been described in detail elsewhere (Page et al., 2021). Ninety-four human neuronal samples with 56 to 70 days in vitro (DIV) belonging to 26 male participants of European ancestry (14 CTR and 12 patients with SCZ) were selected after outlier detection and gene filtering as described ^6^. We further restricted the neuronal samples to pairs present both DIV 56 and 70 (matching by the’SAMPLE ID’ variable in the summarized Experiment object), resulting in a final 27 samples (14 CTR and 13 SCZ) arising from 12 individuals (6 CTR and 6 patients with SCZ). Keeping human genes only, we calculated per gene quantile normalized values from the logRPKM assay. To obtain the gene stability measure, for each gene we computed the across-subjects Kendall’s tau correlation coefficient between quantile-normalized gene expression at DIV56 vs DIV70 [cor.test in R; method = “kendall”,exact=FALSE]).

### Differential Phosphorylation Analysis

SCZ-dependent differential expression log fold change per peptide group was computed within ProteomeDiscover as the mean of SCZ abundance values divided by the mean of CTR abundance values. We compared log2(SCZ/CTR) values between proteins and corresponding phospho-peptides. Within each region, the log2(SCZ/CTR) of phosphorylated peptides was compared to non-phosphorylated peptides using the Wilcoxon rank sum test. Only proteins with corresponding phospho-peptides were considered for this analysis or were shown in the boxplot.

### Gene Set Enrichment Analysis (GSEA)

Ranked gene lists were generated separately for each brain region (CA1, DLPFC, SUB) and data type (protein, phospho-peptide). For each condition, data were filtered by region and type, and the log2(SCZ/CTR) abundance ratio extracted. Proteins and phospho-peptides mapped using Ensembl IDs, missing values were removed and the mean log2(SCZ/CTR) was computed per gene. Conditions with fewer than 50 genes were excluded. Gene set enrichment analysis was performed using the gseGO function from the *clusterProfiler* package with the org.Hs.eg.db annotation database. Analyses were carried out using the Gene Ontology Cellular Compartment ontology, with minimum and maximum gene set sizes set to 15 and 500, respectively, and an adjusted p-value threshold of 0.05.

### Co-expression strength of CLIC sets

CLIC CEM sets were downloaded from https://gene-clic.org/clic/precomputed, comprising 771 sets in total. Mouse MGI.symbol were converted to ensemble.gene.ids using the *getLDS()* function from the *biomart* package. Only genes in the CLIC CEM set that were also in the original ontology list were considered as part of a “CLIC gene set” (CEM.CEM. == “CEM”). A correlation matrix was computed for each dataset through pairwise Pearson correlation of Blom-normalized residual expression. For each CLIC gene set, we cropped the correlation matrix to only genes in the CLIC gene set and to median value of the cropped matrix’s column means to gauge the strength of a CLIC sets connection to itself. This initial co-expression strength value was compared to connectivity to the background, taking into account gene length, GC content and median expression confounders. To this end, we computed a null distribution of co-expression strength, where the rows of the cropped matrix were resampled with genes having a confounder values within the same range of values present in the original CLIC set. A final co-expression strength value was then computed in terms of a z_score in relation to the null:

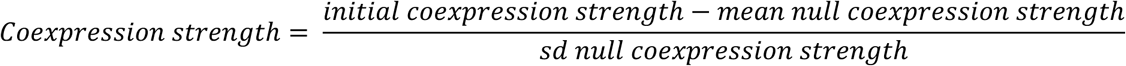

This analysis was performed on LCM expression data (3 different datasets; DLPFC, CA1 and SUB) as well as on bulk tissue data from the Lieber Institute for Brain Development (LIBD) brain repository (3 different datasets; DLPFC, HP and caudate nucleus) ^4,5^. The mean co-expression strength value per gene set was computed within LCM and bulk tissue networks separately. We scaled median co-expression strength values within each dataset (within LCM and within bulk tissue) and compared the scaled values of neuronally relevant sets (see Figure 3A).

### Network identification

We regressed out age, mitochondrial mapping rate, rRNA rate, gene mapping rate from gene and transcript level logTPM data, and regressed out age, total logCPM abundance, and sequencing_batch from protein level logCPM data. We rank-normalized residuals using Blom formula ^72–75^ to limit the impact of deviations from normality in expression data ^76^.

Blom-normalized residuals obtained using the linear models described above were entered as input in the *blockwiseModules* function from the package WGCNA to construct “signed hybrid” network(s), i.e., negative correlation were set to zero and positive correlations were soft-thresholded. We obtained the similarity matrix using Pearson’s R correlation index (*minModuleSize:* 40, *maxBlockSize:* 15,000, *deepSplit:* 4, *mergeCutHeight:* 0.15, *pamStage:* TRUE, *reassignThreshold:* 1e-06). The parameter used for soft-thresholding is the exponent *β* to which the correlation matrix is raised to obtain the adjacency matrix. The standard procedure is to pick the lowest possible β that is high enough to satisfy the scale invariance criterion, defined as the *R*^2^ >.8 in the correlation between original and log-transformed values ^77^. Lower β values are often associated with greater network median connectivity. For each network, we selected the parameter β such that median connectivity was equal across all networks ^6^.

### Gene wise MAGMA

To calculate MAGMA, we used the MAGMA tool v1.09b ^41^, PGC3 summary statistics ^1^ as SNP p-value data, and 1000G European as the reference data file for a European ancestry population to estimate linkage disequilibrium between SNPs. We took the following steps: i) we mapped 1000G SNPs to genes encompassed in each network module (a window of 35-kb upstream and 10-kb downstream of each gene ii) we calculated gene MAGMA Z_scores based on PGC3 SNPs p-values.

### SCZ risk enrichment

SCZ risk genes were those overlapping with 6 varying extension windows around PGC3 GWAS significant SNPs: PGC3 [120 genes], 0 kbp [178 genes] (meaning, genes that overlapped with the index SNP), 20 kbp [299 genes], 50 kbp [456 genes], 200 kbp [1196 genes], 500 kbp [2475 genes] (500 kbp was the maximum extension where enrichments were found significant by ^78^). We assessed the overrepresentation of SCZ risk genes for each module except grey in each network relative to a universe comprised of all genes. We corrected results for multiple comparisons via Bonferroni (number of non-grey modules in each network). Module enrichment of MAGMA was computed using the *gene settest*() from the *limma* package. We labeled as SCZ risk modules those significantly enriched in at least three of the 7 SCZ risk enrichment statistics.

### Jaccard Index analyses

To compute module continuity across expression types, Jaccard Indices (JIs) were computed as the intersection/union of the considered sets. A null distribution for each module pair was computed by resampling each module from their respective universe and recomputing the JI index 10,000 times. A module pair was considered to have significant overlap if the real JI was greater than the max value across the 10,000 permutations, equating to an empirical p_value <.0001.

### Gene-transcript-protein flow

We constructed Sankey plots illustrating the flow of expression across different expression types. Intersection with significant JIs were colored in. Only modules significant for SCZ risk and with significant overlap with a module of another expression type within the same region are shown. The universe was set to the intersection between all network genes in transcript network and all network genes in protein networks. Significant intersections between transcript and protein were evaluated for gene ontologies using the *compareCluster*(fun = enrichGO, ont = “CC”, universe = uni) function from the *clusterProfiler* package, were uni is the intersection between tx and protein universes.

### Interregional circuit-level effects per subject

We defined transcriptomic coupling as the within-subject across-gene correlation (pearson R) of log2(TPM + 1) across all genes expressed in both the regions. Transcriptomic coupling was also computed for CLIC gene sets; each set was taken from CLIC using genes with CEM.CEM. == “CEM” and CEM.CEM. == “CEM +” parameters. Diagnosis effects of transcriptomic coupling were evaluated by comparing differences in the transcriptomic coupling value between CTR and SCZ subjects using a *lm()* taking into account subject-wise confounders of age, mitochondrial mapping rate, rRNA rate, gene mapping rate. The non-scaled pearson R value was used when comparing transcriptomic coupling that was computed across all genes. Since gene set wise transcriptomic coupling is biased within each subject towards the value computed across all genes, we first scaled the values across all gene sets within a subject before comparing across subjects to evaluate diagnosis effects.

### Interregional circuit-level effects of SCZ risk on excitatory synaptic genes

For each gene/protein we computed an across-subject Pearson correlation of one region against another, resulting in correlation matrices of interregional connectivity providing pairwise connectivity values of each and every gene in one region to each and every gene in another region. Pearson correlation was used on Blom-normalized residuals to compute the networks. Interregional co-expression networks were constructed for each region pair, keeping only genes/proteins that are present across both domains and in all three regions (n = 3122).Networks were thresholded to keep only the top 10% of connections.

For each region-pair, we computed the number of retained connections between a specific gene set in one region and a separate gene set of another region. This value was compared to a distribution of connectivity to 1000 null risk sets of the same size and degree distribution to the original set. A null set was adjusted to have a similar degree distribution by replacing each gene in the original set by a random gene that is among the 500 genes with the closest degree values. The Zscoreq95 value describes how many standard deviations the real value is above the 95^th^ quantile of the null, a value above 0 representing an empirical one-sided p <.05. The gene set sizes (*n*) given in the following paragraphs indicate the gene set sizes in the final networks, out of the 3122 remaining genes/proteins.

We evaluated connectivity of pre-synaptic genes (*presynaptic membrane CLIC set, n = 271*) in one region with post-synaptic genes (*dentrite membrane CLIC set, n = 20*) of another region. Each set was taken considering both original ontology genes as well as those found by CLIC to be significantly co-expressed with them (CEM.CEM. == “CEM” and CEM.CEM. == “CEM +”), and removing genes overlapping between sets.

We evaluated connectivity of SCZ risk sets in one region with “excitatory synapse” CLIC genes of another region. We evaluated various types of SCZ risk sets, removing genes found in the “excitatory synapse” set. As a proxy of pure SCZ risk we used the set of all fine-mapped PGC3 genes from Trubetskoy et al., 2022^1^ (*Finemapped.PGC3, n = 81*). Additionally, to proxy a stronger co-expression risk signal, we used a recently published set of genes that are significantly connected to PGC3 genes in DLPFC and/or HP tissue across 3 different consortia, identified by Borcuk et al., 2024 ^50^(*SCZ-connected, n = 74*). Finally, for each region we used the union of all genes in our SCZ risk modules found replicable across transcript and protein WGCNA networks (*REGION_WGCNAreplicated*, those genes described in Figure 4D*, DLPFC n = 12, CA1 n = 29, SUB n = 85*). We then computed connectivity of SCZ risk genes to the CLIC gene set of “excitatory synapse” (*n = 337*). This was taken considering both original ontology genes as well as those found by CLIC to be significantly co-expressed with them (CEM.CEM. == “CEM” and CEM.CEM. == “CEM +”), removing genes found in the corresponding SCZ risk gene set. The null was constructed by keeping the excitatory synaptic genes constant and re-sampling the SCZ risk gene sets.

### Sectioning and sequencing of snRNA-seq of CA1

We performed snRNA-seq on postmortem human brain tissue from the CA1 region for 5 pairs of SZ/Control donors (total of 10 individual samples) using 10x Genomics Chromium Single Cell 3’ Gene Expression technology (v.3). Nuclei were isolated using the “Frankenstein” frozen tissue nuclei isolation protocol developed by Martelotto et. al. ^79^. Briefly, nuclei were made into a suspension by dounce homogenizing 30-40mg frozen tissue in chilled Nuclei EZ Lysis Buffer (MilliporeSigma #NUC101), followed by multiple washes in nuclei wash/resuspension buffer (1x PBS, 1% BSA, 0.2U/μL RNase Inhibitor). The sample was then strained through a 40μm filter, stained with DAPI and AlexaFluor488-conjugated anti-NeuN (MilliporeSigma cat. #MAB377X) to enrich for neuronal nuclei, and sorted on a Beckman MoFlo flow cytometer. We collected 8,500 DAPI+/NeuN+ intact singlet nuclei per sample, sorted directly into 25.1uL reverse transcription reagents from the 10x Genomics Single Cell 3’ Reagents kit (no enzyme).

Libraries were made using the 10x Genomics Chromium Controller platform according to the manufacturer’s instructions. For each cDNA and library sample, quantification and quality control were performed by running a 1:10 dilution of the sample on an Agilent 2100 Bioanalyzer High Sensitivity DNA chip. Libraries were sequenced on a NovaSeq6000 S4 flowcell at Psomagen.

### Pre-processing of snRNA-seq

First, 10x single cell data from all ten samples (five SCZ + five CTR) were individually processed to correct for ambient RNA decontamination using the function *decontX* from R package *celda*. *decontX* uses a bayesian method to estimate and remove contamination in individual cells. To provide correct counts for empty droplets, raw unfiltered matrix files were supplied as background along with the filtered feature matrix files. Counts were rounded to nearest integer before further processing.

Each corrected count matrix was then converted into a Seurat (v4.x) object (min.cells=10, min.features=450). In the first round of filtering, each Seurat object has been filtered individually (2% < nFeature < 98%, 1% < nFeature < 99%). Filtered Seurat objects were temporarily converted to SingleCellExperiment (using *Seurat::as.SingleCellExperiment*) object to add doublet information using *scDblFinder* (nfeatures = 2500) and back to Seurat objects using *Seurat::as.Seurat* keeping both marked singlets and doublets. Next, a single Seurat object has been constructed by merging individual Seurat objects into one. Only genes expressed in at least 400 cells in the merged object were kept. A final round of filtering was performed on the merged Seurat object after inspecting features distribution across individual samples (4% < nFeature < 96%, percent.mt < 8, percent.ribo < 10, Percent.Largest.Gene < 25). Only cells identified as “singlet” were retained afterwards.

After preprocessing, further processing steps from the standard Seurat pipeline were performed on the cleaned object: Normalization (*Seurat::NormalizeData*(normalization.method = “LogNormalize”, scale.factor = 10000))-> Finding variables features (*Seurat::FindVariableFeatures*(selection.method = “vst”, nfeatures = 2000))-> Scaling on all genes (*Seurat::ScaleData*)-> PCA on all features (*Seurat::runPCA*). To minimize the batch effects among samples, we employed *harmony::RunHarmony* integration function with default parameters and original sample names as the batch covariate. Downstream dimensional reduction methods like UMAP and tSNE used harmony embeddings instead of default PCA embeddings.

### CellChat

Inter-cell type ligand-receptor interactions were identified using the package *CellChat* ^51^. CellChat objects were run separately using either only CTR or only SCZ subjects. Over-expressed interactions were first identified using the *identifyOverExpressedGenes* then *identifyOverExpressedInteractions,* using default parameters. This defines over-expressed ligand-receptor interactions if either ligand or receptor is over-expressed in either of the two interacting cell types. Only these over-expressed ligand-receptor interactions are considered in further steps. A communication probability was then computed for each interaction using *computeCommunProb*(cellchat, type = “truncatedMean”, trim = 0.1), and cell types with fewer than 10 cells were removed using *filterCommunication*(cellchat, min.cells = 10). Celltypes were grouped based on similar ligand inputs using *identifyCommunicationPatterns*(cellchat_real, pattern = “incoming”, k = 4). Differential numbers of interactions were visualized using *netVisual_diffInteraction*(cellchat, weight.scale = T, measure = “count.merged”, label.edge = T). SCZ-dependent overexpressed genes were computed per cell type using “net = *identifyOverExpressedGenes*(cellchat, only.pos = FALSE, thresh.pc = 0.1, thresh.fc = 0.05,thresh.p = 0.05, group.DE.combined = FALSE)”. Significantly differentially expressed genes were mapped onto significant ligand-receptor interactions using *netMappingDEG*. Up-regulated genes found only in SCZ-dependent interactions were identified using *subsetCommunication*(cellchat, net = net, datasets = “SCZ”,ligand.logFC = 0.05, receptor.logFC = NULL). Down-regulated genes found only in CTR-dependent interactions were identified using *subsetCommunication*(cellchat, net = net, datasets = “CTR”,ligand.logFC =-0.05, receptor.logFC = NULL). We visualized canonical interneuronal inputs to Excit_A from inhibitory clusters using *netVisual_bubble*(cellchat, sources.use =c(“Inhib_B”, “Inhib_C”, “Inhib_D”), targets.use = “Excit_A”, signaling = c(“NPY”,“VIP”,“SOMATOSTATIN”, “OPIOID”)).

### AUCell gene set enrichments of celltypes

To identify celltype clusters that are enriched for schizophrenia associated gene-sets, we employed the AUcell enriched method. The AUCell algorithm calculates enrichment scores by determining the proportional area under the curve of gene expression rankings for each gene set in individual cells. In order to employ this method we specifically chose a package called Escape^80^ which is a bridging package to facilitate gene set enrichment analysis in snRNA-seq data. The runEscape() function was used to calculate the enrichment scores with AUC method (method = “AUCell”), and the minimum number of gene set genes to be detected in a celltype to compute a score as 10 (min.size = 10). We evaluated enrichment of the *Finemapped PGC3* and *REGION_WGCNAreplicated* SCZ risk gene sets as described in the “Interregional circuit-level effects of SCZ risk on excitatory synaptic genes” methods section. Additionally, we used a set of genes that are significantly connected to PGC3 genes in the HP tissue across 2 different consortia, identified by Borcuk et al., 2024 ^50^(*SCZ-connected_HP*).

## Acknowledgements

We would like to thank Nicola Pedreschi for aid in development of the interregional circuit-level analyses, and Nina Rajpurohit for aid in writing the snRNAseq methods.

