## Supplementary Information for "Transcriptomics and proteomics of projection neurons in a circuit linking hippocampus with dorsolateral prefrontal cortex in the human brain"

### Supplementary methods

#### Identifying a neuronal CLIC community

A p<.001 significance value was associated to the Jaccard index (JI) of all possible hypothetical pairs of CLIC sets with sizes of 10,25,50,100,250,500,1000,2500,5000, or 7500. This was run by resampling the two hypothetical CLIC sets from the universe of all CLIC genes 10000 times and assigning the $5^{th}$ highest permuted JI to p<.001. These values were used to construct a model to predict the JI value associated with p<.001 based on the sizes of the two CLIC sets, using lm(p001_JI ~ GS1_size*GS2_size ). Real CLIC set pairs with a JI greater than their predicted null JI value were considered to have significant gene overlap. A graph was created with nodes as CLIC sets and edges between sets with significant gene overlap, and an edge weight equal to the pair's real JI minus its predicted p001_JI. The weighted graph was clustered using the Louvain algorithm, resolution = 1 (Figure S1A). Within each community overlapping genes were considered as those that appeared in a cluster 4 times as frequently as outside of a cluster, (i.e. log2(FC) >= 2). This identified 4 primary communities with 3 having clear GO enrichments of within community overlapping genes related to gene transcription or translation (Figure S1B, three right most), and another community having mixed ontologies. This mixed community was separated as a subgraph and re-clustered using the Louvain algorithm, resolution = 1 (Figure S1A, bottom). This identified 5 communities with clear ontologies (Figure S1B, five left most), among which one community, termed "Neuronal", contained only neuronally relevant ontologies (Figure S1B, most left). The CLIC set names within this community also show clear over-representation of neuronal terms (Figure S1C).

### Supplementary figures

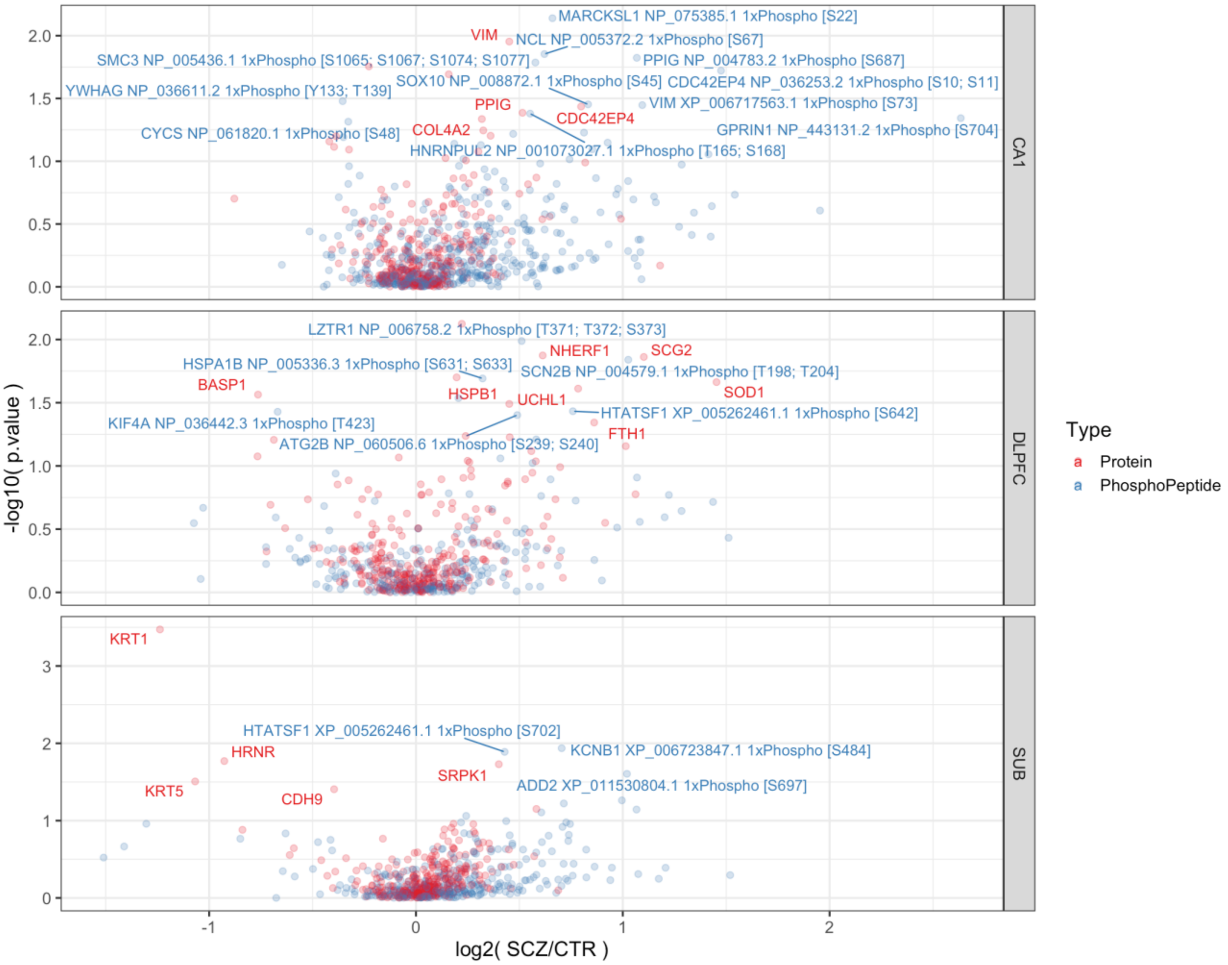


**Supplementary Figure S1. Volcano plot of protein and phospho-peptide SCZ-dependent differential abundance**
Volcano plots depicting SCZ-dependent differential abundances for proteins and their phospho-peptides separately (differentiated by color) and per region (differentiated by panel). Fold changes are shown on the x-axis and nominal significance is shown on the y-axis. Only proteins with corresponding phospho-peptides present are plotted here. Phospho-peptides and proteins with nominal p values less than .05 are labelled. Phospho-peptides labels include the master protein accession id and phosphorylation site.

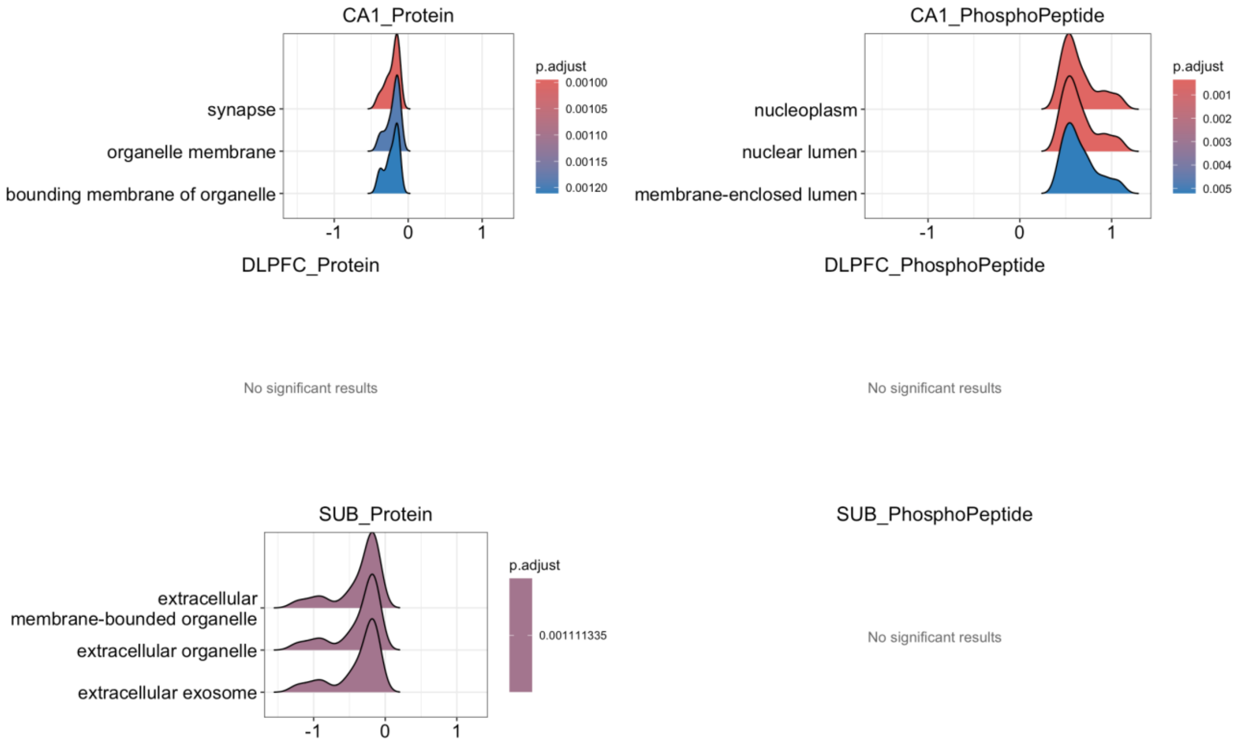


**Supplementary Figure S2. Geneset enrichment of SCZ-dependent differential abundance**

We used the function gseGO to cellular compartment ontologies that contain genes with log2(SCZ/CTR) values significantly higher (or lower) than the background. Peptides and proteins were annotated to genes, and the mean log2(SCZ/CTR) value across phospho peptides or proteins was attributed to each gene. Ridgeplots depict the distribution of log2(SCZ/CTR) values (y-axis) for significant ontology genesets (x-axis). Each panel represents a combination of region (CA1 top, DLPFC middle, SUB bottom) and measure type (protein on the left, phospho-peptide on the right).

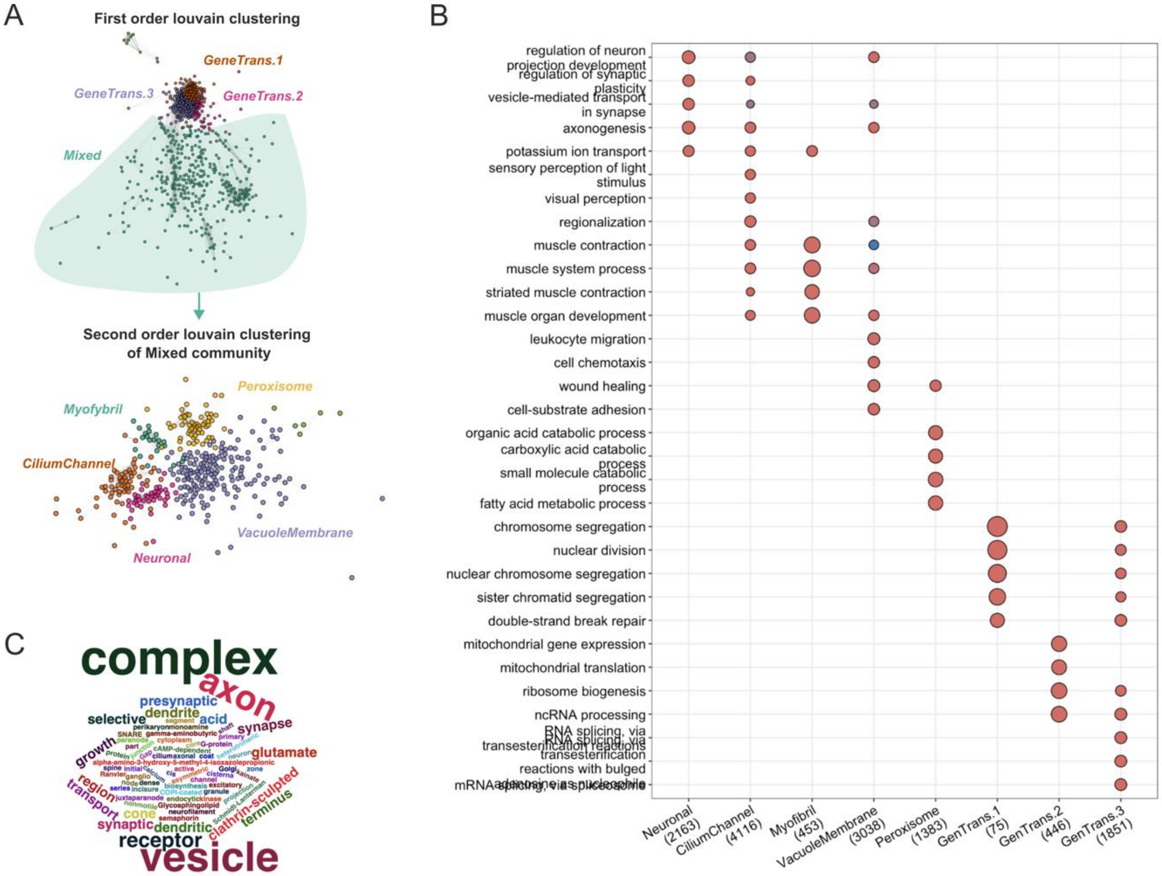


**Supplementary Figure S3. CLIC set communities**

A) CLIC sets were clustered based on whether they have overlapping genes

B) Communities were assigned identities based on GO enrichment of overlapping genes. This identified a neuronal CLIC community.

C) The CLIC set names within the neuronal community show strong representation of neuronal terms.

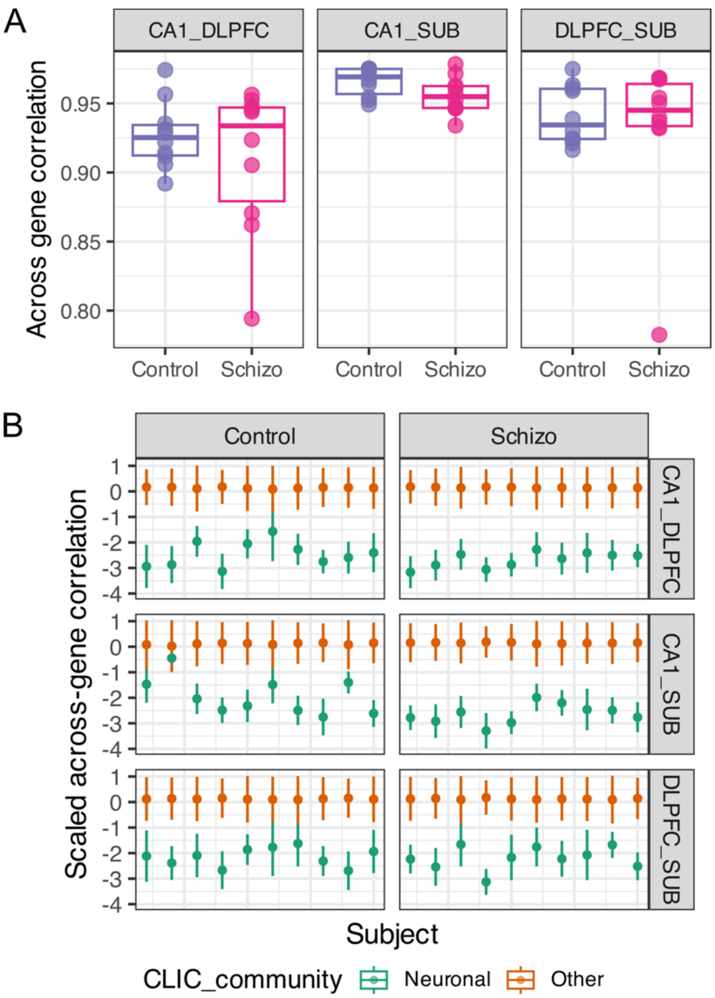


**Supplementary Figure S4. Additional results related to interregional circuit-level effects per subject**

A) To evaluate transcriptomic coupling between region pairs, we computed the across-gene correlation strength (Pearson R) per subject per region pair, computing across the union of all CLIC genes. On the y-axis is the across gene correlation strength. Points are subjects, colored and spread on the x-axis by diagnosis.

B) We computed the across-gene correlation strength for each CLIC gene set, each time across only the genes within the set. The values scaled within each subject, found on the y-axis. Points and error bars representing neuronal gene sets containing neuronally relevant key terms are colored in green, and all other gene sets are colored in grey. The points depict the median, and the bars depict the sd. Communities were assigned identities based on GO enrichment of overlapping genes. This identified a neuronal CLIC community.
